# Recruitment of SH3P2–DRP1A by HYCCIN2 Drives Membrane Tubulation in Arabidopsis Embryonic Cell Plate Formation

**DOI:** 10.1101/2024.09.06.611148

**Authors:** Ya-Wen Hsu, Chine-Ta Juan, Cian-Ling Guo, Huei-Jing Wang, Guang-Yuh Jauh

## Abstract

Cytokinesis marks the culmination of cell division when the cytoplasm undergoes division to yield two daughter cells. This intricate process encompasses various biological phenomena, including the organization of the cytoskeleton and the dynamics of vesicles. The development of the cell plate aligns with alterations in the cytoskeleton structure and vesicles derived from the *trans*-Golgi, ultimately resulting in the formation of the planar cell plate. Nevertheless, the coordination of these processes in plants remains to be determined. Here, we introduce HYCCIN-CONTAINING2 (HYC2) as a pivotal cytoskeleton cross-linking protein that accumulates at phragmoplasts and plays a crucial role in cell plate formation. A genetic study involving the depletion of HYC2 post-anaphase in Arabidopsis revealed HYC2’s function in cell plate formation. HYC2 interacted with dynamin-related protein 1A (DRP1A) and SH3 domain-containing protein 2 (SH3P2), essential for membrane tubulation during cell plate formation. The recruitment of SH3P2 and DRP1A to the cell plate and phragmoplast organization was compromised in homozygous *hyc2-2* mutant globular embryos. Our results shed light on the cytoskeletal function of HYC2 in assembling the cell plate, potentially by guiding vesicles containing SH3P2–DRP1A to the planar cell plate.

**Significance statement:** During plant cell division, a new membrane is constructed at the division plane, and vesicles fuse to form a transient compartment known as the cell plate. This structure undergoes stabilization via membrane remodeling events, including tubulation, to generate robust membrane intermediates. Apart from DRP1A and SH3P2, our understanding of other protein components involved in membrane remodeling still needs to be completed. HYC2 is a plant counterpart to the human Hyccin protein involved in neuronal myelination and displaying bundling capabilities for microtubules and microfilaments. Our study demonstrates that HYC2 is crucial in recruiting proteins during membrane tubulation in the cell plate formation in developing Arabidopsis embryos. These findings underscore the molecular function and vital role of the Hyccin protein in cell plate formation.

**One Sentence Summary:** Plant HYC2 displays bundling capabilities for microtubules and microfilaments to recruit DRP1A and SH3P2 during membrane tubulation in the cell plate formation in developing Arabidopsis embryos.

## Introduction

Within the eukaryotic cell division cycle, cytokinesis is the conclusive step in physically segregating the emerging daughter cells. In metazoans and fungi, this process initiates from the periphery. It progresses toward the center via the formation of a contractile actomyosin ring, leading to the constriction of the cell along the division plane. However, flowering plants, lacking myosin II genes and the formation of a contractile ring, have evolved an alternative cytokinesis strategy. The strategy involves the homotypic fusion of secretory vesicles to the partitioning membrane along dynamic microtubule tracks in the phragmoplast to construct the cell plate (Rasmussen et al., 2011; Jurgens et al., 2015). Vesicles derived from the *trans*-Golgi network undergo fusion, resulting in hourglass-shaped vesicle intermediates. These intermediates are then integrated into the cell equator, forming a tubulo-vesicular network, a tubular network, and a planar fenestrated sheet (PFS). Concurrently, cell wall polymers such as callose gradually accumulate within the tubulo-vesicular network, thus helping to stabilize its structure up to the PFS. Ultimately, the expanding PFS fuses with the mother cell wall (Smertenko et al., 2018).

Following anaphase, the phragmoplast emerges as a scaffold crucial for orchestrating the cell plate assembly and forming a new cell wall during cytokinesis. Comprising a complex arrangement of microtubules and microfilaments, the phragmoplast plays a pivotal role in facilitating the movement of secreted vesicles and positioning cell wall components for incorporation into the cell plates (Jurgens et al., 2015; Smertenko et al., 2018). In telophase, the phragmoplast manifests as two sets of antiparallel microtubules that converge and overlap at the midline. Materials containing secreted vesicles, essential for constructing the cell plate, traverse along these microtubules and merge to give rise to the cell plate. Consequently, proteins that modulate cytoskeletal dynamics, such as microtubule-associated proteins (MAPs) or actin-binding proteins (ABPs), play a fundamental role in the division and development of plant cells.

In embryo development, only a few cytoskeletal proteins have been identified, including the microtubule-severing protein KATANIN 1 (Luptovciak et al., 2017), microtubule-associated protein 65-3 (MAP65-3; Caillaud et al., 2008; Ho et al., 2011), HINKEL (Strompen et al., 2002), kinesin-related protein 125c (AtKRP125c; Bannigan et al., 2007), and RUNKEL (Krupnova et al., 2009). These cytoskeletal proteins regulate the organization of the phragmoplast microtubule array but not the trafficking of regulatory proteins to the cell plate. The coordination between cytoskeleton proteins and vesicle targeting during cell plate formation remains elusive, as indicated by the use of cytoskeletal inhibitors suggesting an interaction between the two cytoskeletal tracks whereby disrupting one type of filament affects the stability of the other (Maeda et al., 2020).

Hypomyelination and congenital cataract (HCC) was initially identified as a human white-matter disorder characterized by the insufficient myelination of both the central and peripheral nervous systems, thus resulting in progressive neurological decline and the presence of congenital cataracts (Zara et al., 2006). HCC is attributed to a deficiency in the Hyccin protein, a novel neuronal protein primarily expressed in the central nervous system and localized explicitly in neurons rather than myelinating cells (Zara et al., 2006; Ugur and Tolun, 2008; Gazzerro et al., 2012; Traverso et al., 2013). The plasma membrane-anchored phosphatidylinositol 4-kinase scaffolding complex comprises Hyccin, PI4KIIIα, and its adaptors tetratricopeptide repeat domain 7 (TTC7) and pho eighty-five requiring 3 (EFR3) (Baskin et al., 2016). Dysfunction of the Hyccin protein in fibroblasts from HCC patients and the oligodendroglial lineage of knockout mice feature reduced levels of PI4KIIIα, TTC7A/B, and EFR3A, which indicates the role of Hyccin in regulating PtdIns(4)P production. In the Arabidopsis genome, two *HYCCIN-CONTAINING* genes, *HYC1* and *HYC2*, encode proteins as components of the PI4Kα1 complex. HYC1 is essential for the male gametophyte, whereas HYC2 is crucial for embryogenesis (Noack et al., 2022). However, the underlying molecular mechanism of HYC2 in embryogenesis remains to be elucidated.

In a search to identify genes crucial for embryogenesis in Arabidopsis, we obtained the *hyc2* mutant, which features embryonic arrest at the late globular stage, forming aborted seeds. Biochemical analysis revealed the molecular role of HYC2 as a protein that bundles microfilaments and microtubules. HYC2 protein accumulates in the phragmoplast and is involved in cell plate formation, suggesting its involvement with microtubules and microfilaments. Additionally, we observed an interaction between HYC2 and the SH3P2– DRP1A complex, recognized for its role in membrane tubulation in cell plate formation. In *hyc2-2/-* globular embryos, signals from SH3P2 and DRP1A were absent at the developing cell plate. Hence, the essential function of the HYC2 protein in the cytoskeleton is crucial for recruiting the SH3P2–DRP1A complex to the cell plate in Arabidopsis.

## Results

### Disruption of HYC2 results in the arrest of globular embryos and the formation of cell wall stubs in Arabidopsis embryos

As the results found by Noack *et al*., 2022, we observed seed abortion phenotype in the heterozygous mutants of *hyc2-2* and *hyc2-3* (Figure S1A), and a quarter of the embryo was arrested at the globular stage when most had reached the heart stage in individual *hyc2/+* siliques (Figure S1B). Genotyping of grown seedlings from heterozygous *hyc2/+* progenies revealed a 1:2 ratio of wild type to heterozygotes, indicating defective embryogenesis (Table S1). The primary amino acid sequence of HYC2 shares 35.8% identity with the Hyccin domain (pfam0970), which is present in eukaryotic Bilateria and plants but absent in unicellular yeast. Protein alignment demonstrated high conservation of the Hyccin domain, with divergence in the C-terminal region (Figure S2). *HYC2* was ubiquitously expressed in all tissues, with the highest expression in siliques (Figure S3A). The seed abortion phenotype of *hyc2-2* was successfully rescued by its transgene *HYC2p::HYC2-2XGFP* (Figure S1A). Genetic and complementation analyses suggested that the arrested globular seeds in *hyc2-2/+* siliques are *hyc2-2* homozygotes (*hyc2-2*). The seed abortion phenotype of *hyc2-2* could be rescued by full-length HYC2, HYC2^1-417^, and HYC2^1-405^, but not HYC2^1-398^ (Figure S3B). JPred secondary structure prediction (Drozdetskiy et al., 2015) indicated that the penultimate helix of HYC2 (375-395 aa) is crucial for complementation (Figure S2A). These findings underscore the essential role of HYC2 in proper embryo development in Arabidopsis.

Because of arrested globular embryos in homozygous *hyc2-2/-*, we used SCRI Renaissance 2200 (SCRI), a dye for cell wall staining, to delineate the developing embryo outline (Smith and Long, 2010). SCRI staining revealed that cell organization was well coordinated in the wild type, and cell wall development proceeded normally (Figure 1A). However, in the *hyc2-2/-* embryo, cell wall formation exhibited irregularities and occasional discontinuities. Some embryonic cells featured a noticeable cell wall discontinuation phenotype (indicated by arrowheads in Figure 1A), reminiscent of observations in cytokinesis-deficient mutants (Sollner et al., 2002). Moreover, the cell number in the *hyc2-2/-* embryo exceeded that of the wild type at the globular stage and approached that of the wild type at the transition stage (Figure 1A, Table S2). We also investigated potential cytokinesis defects in *hyc2-2/-* embryos using ultrastructural observation (transmission electron microscopy [TEM] images in Figure 1B). In the wild-type embryo, the cell wall appeared smooth and compact, whereas, in the *hyc2-2/-*embryo, the cell wall featured looseness, protrusions, and occasional failure to fuse with the parental cell wall. The SCRI and TEM results, portraying discontinuous cell plates with undulating membrane sacs linked to incomplete cell plate formation, underscore the crucial role of HYC2 during cell plate formation in Arabidopsis.

**Figure 1.**
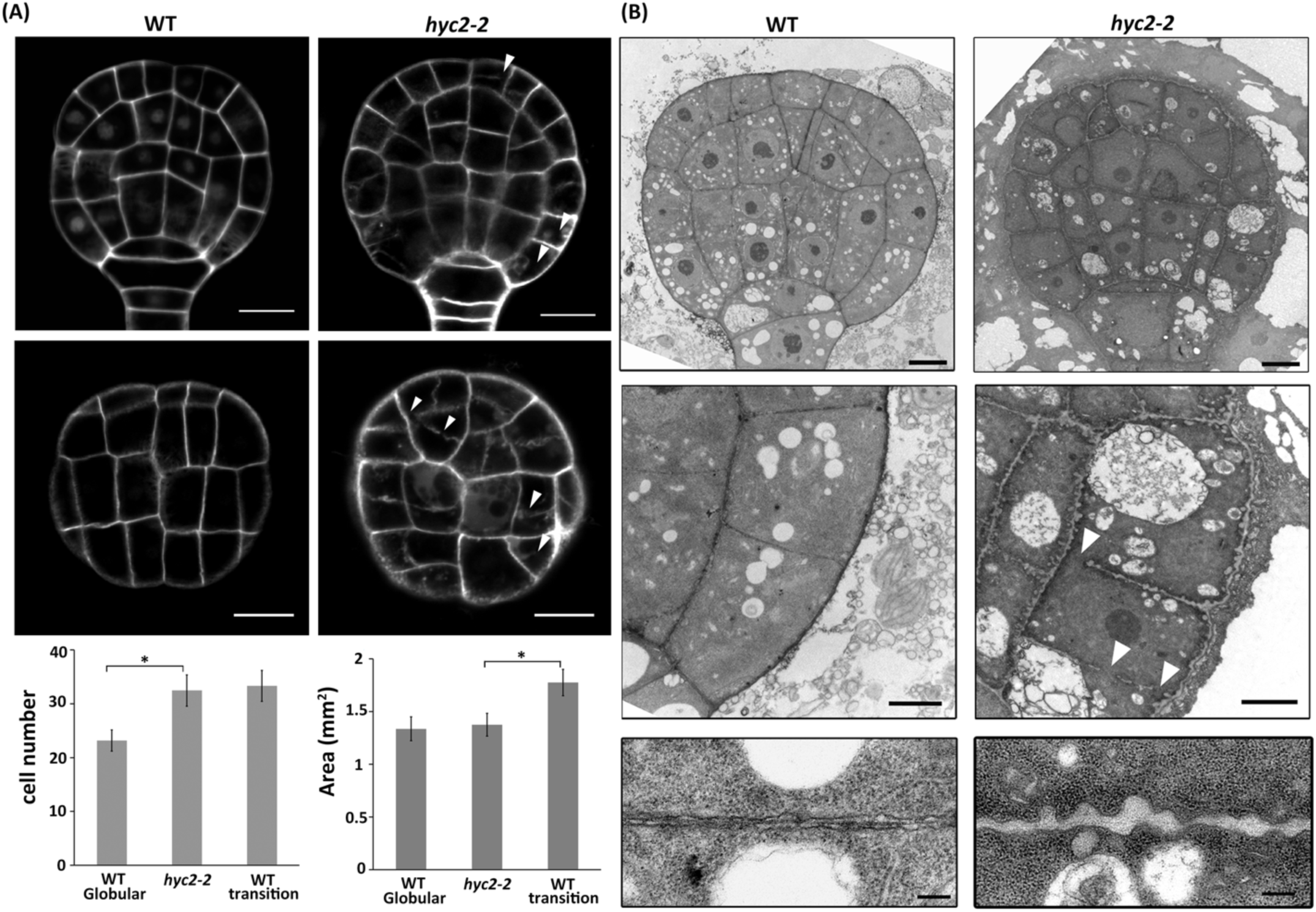
Cytokinesis defects in globular embryos of *hyc2-2/-* mutants. (A) SCRI-stained globular embryo in the WT and *hyc2-2*/-. Homozygous *hyc2-2/-* globular embryos exhibit a leaky cell wall phenotype, indicated by white arrowheads—bar=10 μm. The cell number and cell area of globular embryos in the WT and *hyc2-2/-* were analyzed using ImageJ (n=20 and n=17 for WT and *hyc2-2/-*; Data are mean±SEM, *p*<0.0001, two-tailed, Student *t* test, n=3 independent experiments). (B) Ultrastructural observation of globular embryos in the WT and homozygous *hyc2-2/-*. White arrowheads indicate the leaky cell wall in *hyc2-2/-* embryos; bar = 5 μm, 2 μm, and 0.2 μm from the top to the bottom of the panel.

### HYC2 is a component of the PI4Kα1 complex and is critical for the completion of cell plate formation

HYC2 interacts with No Pollen Germination (NPG) proteins and serves as a component of the PI4Kα1 complex (Noack et al., 2022). To elucidate HYC2’s role within the PI4Kα1 complex, we examined PI4P signals using the fluorescent signal of theYFP-PH_FAPP1_ (Vermeer et al., 2009). *In planta*, PI4P signals primarily accumulated at the plasma membrane in wild-type embryos from the globular to the linear stage (Figure S4A). In contrast, in *hyc2-2* globular embryos, the PI4P signal decreased at the plasma membrane and began accumulating in the cytosol (Figure S4A), which suggests HYC2’s involvement in recruiting PI4P to the plasma membrane. In dividing tobacco BY-2 cells, YFP-PH_FAPP1_ labeled the growing cell plates after detecting FM4-64 signals (Vermeer et al., 2009). Similarly, in Arabidopsis root cells, PI4P signals were gathered at the expanding cell plate, already marked by HYC2 signals (Figure S4B). HYC2 signals appeared in the cell plate before the PI4P signals, implying HYC2’s role in cell plate formation.

To gain insights into the cellular role of HYC2 in cell plate formation, we examined its localization in dividing cells. In Arabidopsis embryogenesis and root cells, HYC2 was found in the cytosol and plasma membrane (Figure 2). The fluorescent signals of HYC2-2XGFP, driven by the native promoter, exhibited robust HYC2 expression after the 8-cell embryo stage (Figure 2A). Additionally, time-lapse fluorescence microscopy revealed HYC2 accumulation at the cell plate during the division of root cells (Figure 2B). HYC2-GFP appeared in the midsection of the cell, where cell plate formation initiates, before labeling the emerging cell plate by FM4-64 signals (Dhonukshe et al., 2006). It gradually surrounded the FM4-64 signals until the cell plate formation was completed. The organization of the cytoskeleton, including microtubules and microfilaments, plays a regulatory role in cell division. To explore the connection between HYC2 and the cytoskeleton in cell plate formation, we used immunofluorescence analysis at different stages of mitosis. HYC2 was co-localized with the microtubule array after metaphase (Figure S5A) and with F-actin signals after telophase (Figure S5B). Hence, HYC2’s function is crucial in cell plate formation.

**Figure 2.**
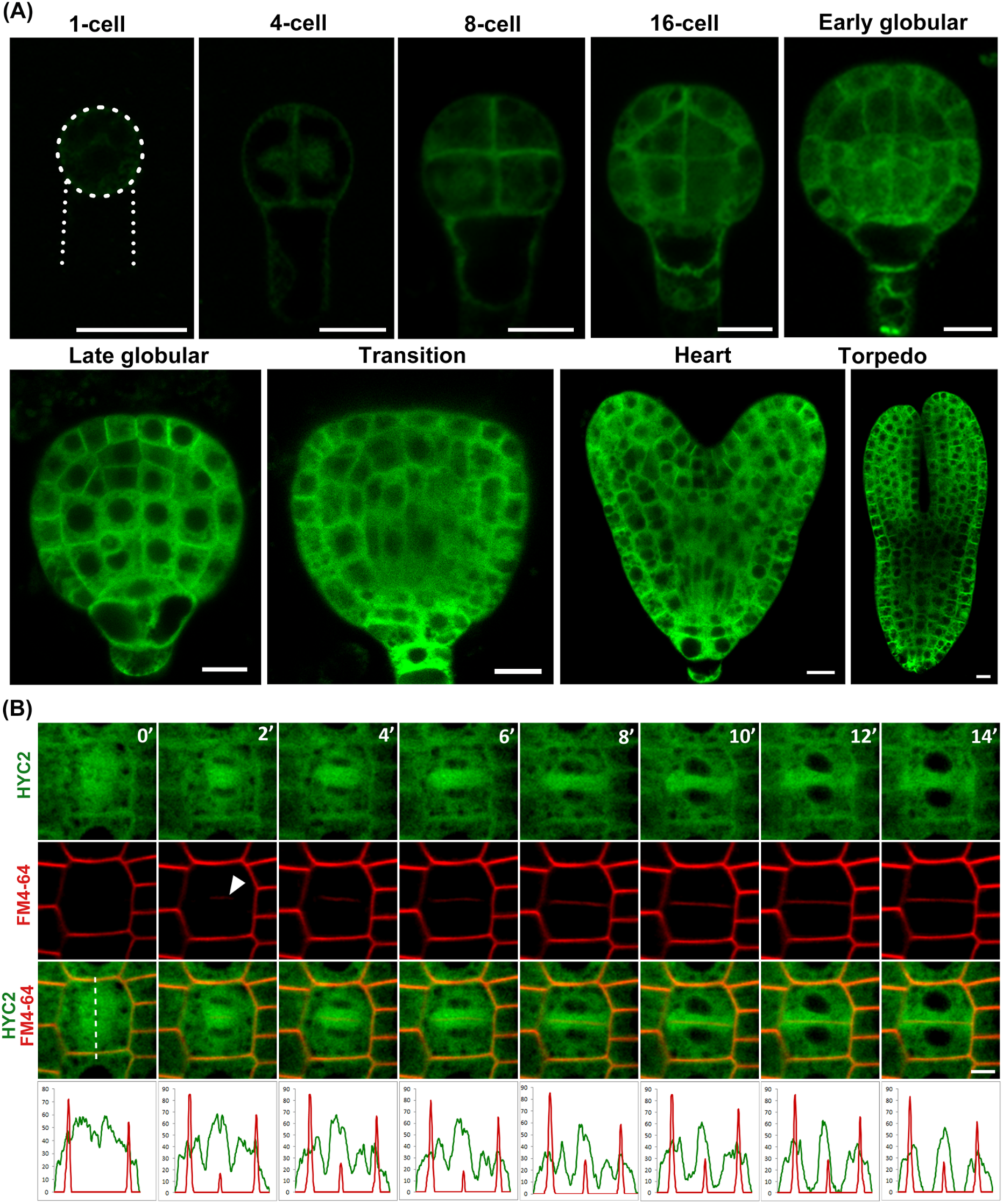
Subcellular localization of HYC2 in Arabidopsis embryos and root tips. (A) HYC2 is localized in developing embryos using the HYC2-2XGFP fusion driven by the HYC2 native promoter. HYC2-2XGFP signals are observed from 1-cell embryos to torpedo-stage embryos, with the outline of a 1-cell embryo indicated by a white dotted line. Bar = 10 μm. (B) Time-lapse observation of root tip cells in 7-day-old seedlings expressing *HYC2pro::HYC2-2XGFP*. Images were captured at 3-minute intervals. The bottom panel illustrates the quantified fluorescence intensity of HYC2-2XGFP (green) and FM4-64 (red) corresponding to the signals from the relative position of the dashed line at each time point. The white arrowhead marks the first observation of the emerging cell plate. Bar = 5 μm.

To further investigate the role of HYC2 in cell plate formation, we rescued *hyc2-2/+* embryos by using GFP-tagged HYC2 fused with the N-terminal region of *CYCB1;2*, containing the destruction box (D box) responsible for protein degradation in anaphase (Criqui et al., 2000). HYC2-CYCB1;2 was progressively degraded concomitant with the emergence of the cell plate marked by FM4-64 (Figure 3A). As compared with our previous findings indicating the rescue of the seed-aborted phenotype of *hyc2-2/+* by *HYC2pro::HYC2-2XGFP* (Figure 1A), 26.8% of seeds were arrested at the globular stage in *HYC2pro::HYC2-CYCB1;2-2XGFP*/*hyc2-2/+* and in *hyc2-2/+* (Figure 3B). This result underscores the need for the presence and function of HYC2 after the onset of anaphase for successful cell plate formation and embryogenesis in Arabidopsis.

**Figure 3.**
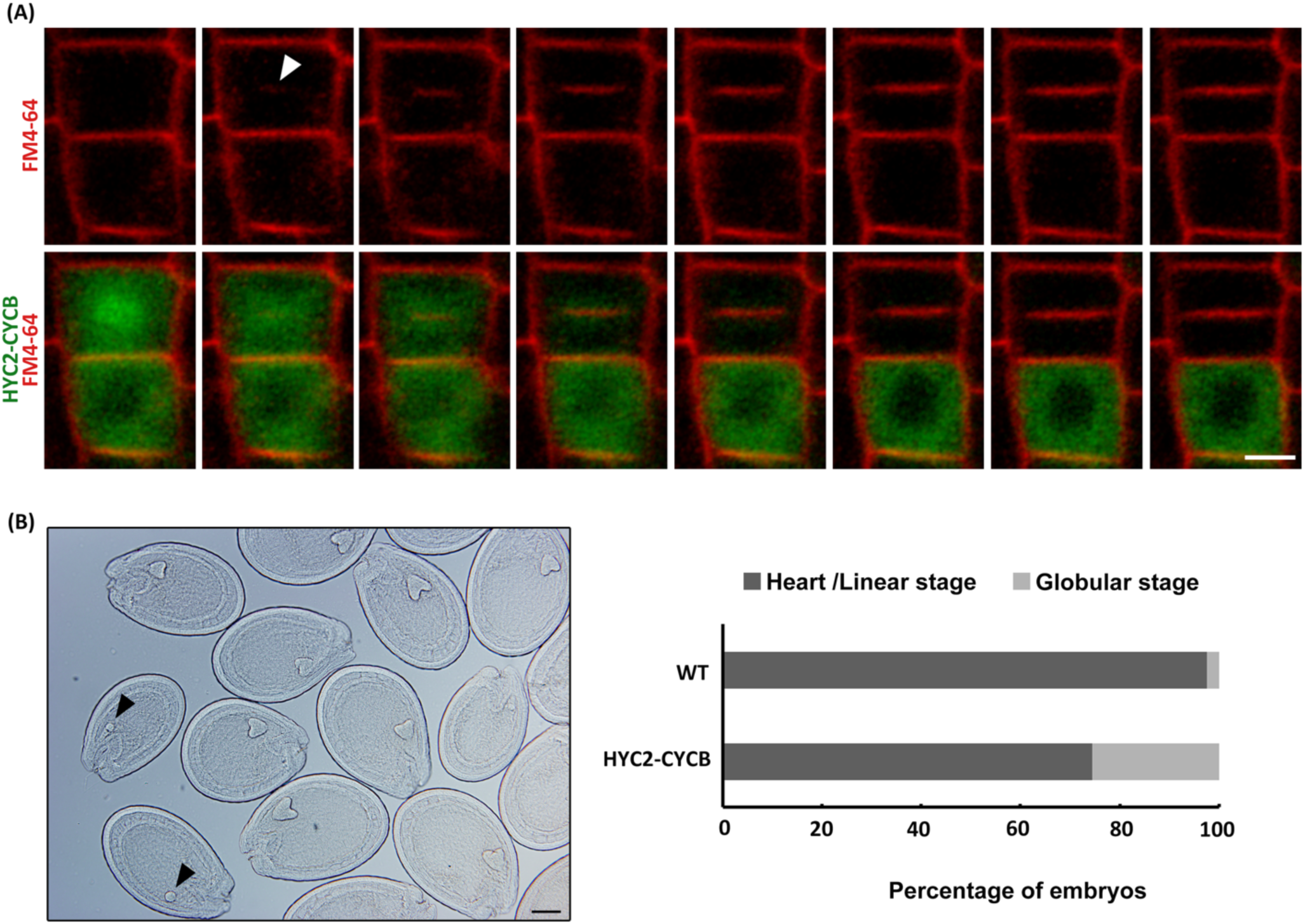
HYC2 plays an essential function during the post-anaphase stage of embryogenesis. (A) Time-lapse observations of root tip cells in 7-day-old seedlings expressing *HYC2pro::HYC2-CYCB1;2-2XGFP*. White arrowheads mark the emergence of the cell plate. Images were captured at intervals of 3 min. (B) Color clearance of the developing silique from *HYC2pro::HYC2-CYCB1;2-2XGFP*/*hyc2-2/+*. The plot showed the percentage of embryo developmental stage in the developing siliques, and approximately 26.8% of embryos were arrested at the globular stage, and the remaining embryos stopped at the heart stage (n=121 and n=227 for WT and *HYC2pro::HYC2-CYCB1;2-2XGFP*/*hyc2-2/+*, respectively). *p* > 0.05 (χ2 test, WT/HZ: HM=3:1, n=3 independent experiments). Globular embryos in the siliques of *HYC2pro::HYC2-CYCB1;2-2XGFP*/*hyc2-2/+* are indicated by black arrowheads.

### HYC2 exhibits binding and bundling properties for both microtubules and microfilaments

To elucidate the biological function of HYC2, we tried to identify the potential HYC2-interacting proteins by co-IP and liquid chromatography-tandem mass spectrometry (LC-MSMS) analyses. Numerous tubulin and actin proteins were co-immunoprecipitated with HYC2 (Table S3), and their interactions were confirmed by immunoblotting assays (Figure 4A). To determine the roles of HYC2 in tubulin and microtubule binding ability, microscale thermophoresis (MST) and co-sedimentation analysis (Figure 4B) were further used. The results illustrate the direct interaction of recombinant HYC2 with tubulin, thereby suggesting its role as a tubulin-binding protein. Low-speed co-sedimentation assay between microtubule and HYC2 revealed the microtubule bundling activity of HYC2 (Figure 4D upper panel). *In vitro* bundling experiments clearly showed clustered microtubule accumulation in the presence of HYC2, and these HYC2-bundled microtubules were resistant to cold treatment-induced disassembly (Figure 4D lower panel).

**Figure 4.**
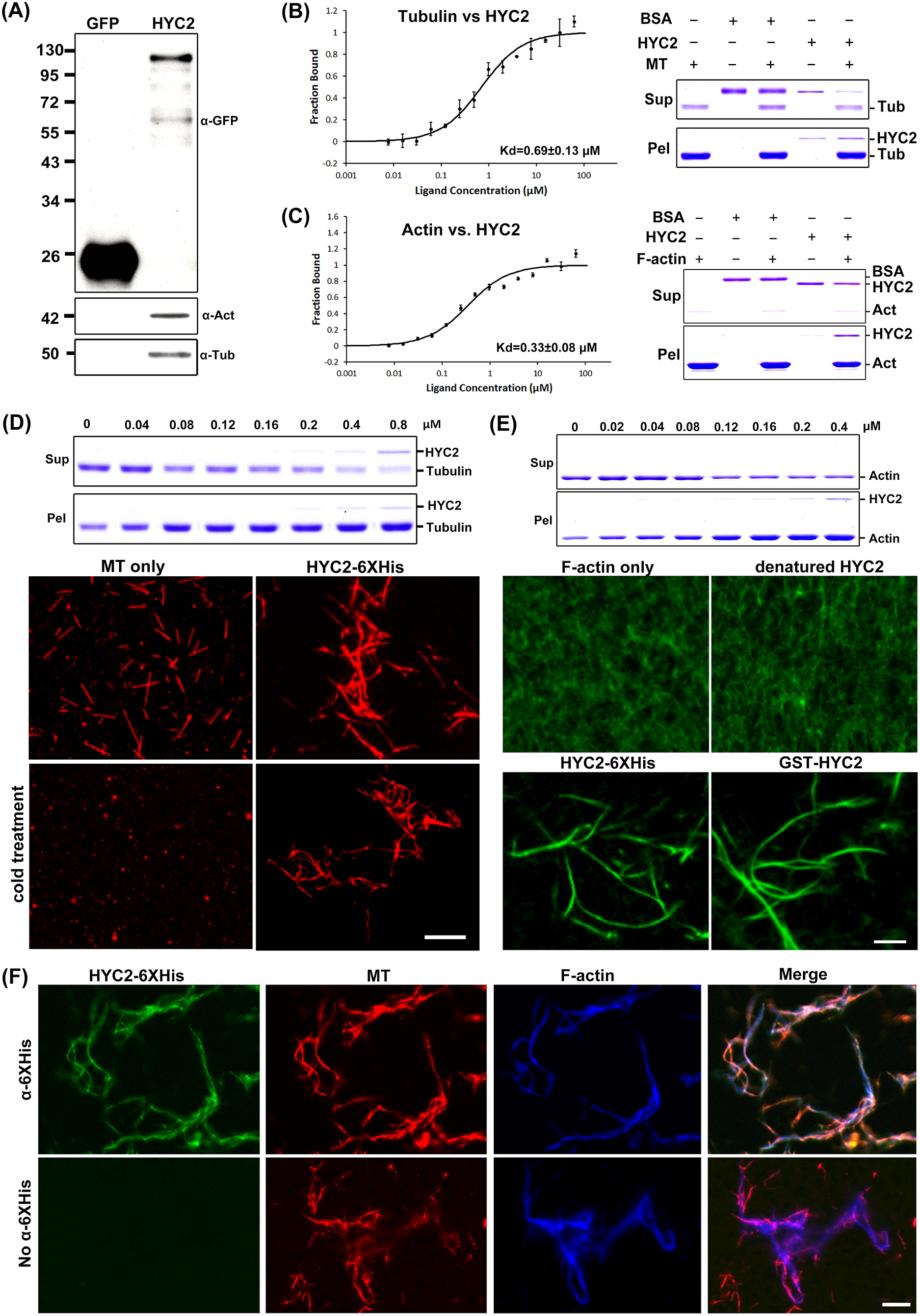
HYC2 functions as the microtubule and microfilament binding and bundling protein. (A) Immunoblot analysis of anti-GFP immunoprecipitates from 7-day-old seedlings expressing HYC2-2XGFP or GFP alone. The indicated antibodies were used for analysis. (B) MST measurements and high-speed co-sedimentation assays showed the interaction of HYC2 with tubulin and microtubules. For MST measurements in the left panel, a fixed concentration of 34 nM Alexa-647 NHS-labelled tubulin was titrated with 62.5 μM to 7.6 nM HYC2. An equilibrium binding constant of 0.69±0.13 μM was determined by fitting a hyperbola to the data. Soluble BSA was chosen as a negative control for *in vitro* microtubule-binding assay using high-speed co-sedimentation in the right panel. Pel: pellet fraction; Sup: supernatant fraction. (C) MST measurements and high-speed co-sedimentation assay showed the interaction of HYC2 with actin and F-actin. For MST measurements in the right panel, a fixed concentration of 38.8 nM Alexa-647-labelled actin was titrated with 62.5 μM to 7.6 nM HYC2. An equilibrium binding constant of 0.33±0.08 μM was determined by fitting a hyperbola to the data. Soluble BSA was chosen as a negative control for in vitro F-actin-binding assay using high-speed co-sedimentation in the right panel. Pel: pellet fraction; Sup: supernatant fraction. (D) Microtubule bundling activity of HYC2 was determined by low-speed co-sedimentation assay and fluorescence microscopy. A low-speed co-sedimentation assay for *in vitro* microtubule-bundling activity was used in the upper panel. The HYC2 protein level is indicated above the gel. Pel: pellet fraction; Sup: supernatant fraction. In the bottom panel, microtubule bundles induced by recombinant HYC2-6XHis protein were revealed by fluorescence microscopy. Rhodamine-labeled microtubules (red) were co-incubated with HYC2 recombinant protein under cold treatment. Bar = 5 μm. (E) The low-speed co-sedimentation assay and fluorescence microscopy demonstrated the F-actin bundling activity of HYC2. Low-speed co-sedimentation assay for *in vitro* F-actin-bundling activity under 10 μM Ca^2+^ condition was shown in the upper panel with the indicated HYC2 protein levels above the gel. Pel: pellet fraction; Sup: supernatant fraction. In the bottom panel, F-actin bundles labeled by Alexa 488-phalloidin (green) induced by recombinant HYC2-6XHis and GST-HYC2 proteins were observed using fluorescence microscopy. Bar = 5 μm. (F) Immunofluorescence demonstrated the simultaneous co-localization of HYC2 with microtubules and microfilament bundles. The anti-His antibody (GFP, upper panel) recognized HYC2-6XHis signals but not the control without the anti-His antibody (lower panel). HYC2 co-localized well with bundles composed of rhodamine-labeled microtubules (red) and Alexa 633-phalloidin-stained F-actin (blue). Bar = 5 μm.

The MST and co-sedimentation assays were also applied to demonstrate the direct interaction of recombinant HYC2 with actin and F-actin (Figure 4C). The F-actin co-sedimentation assay showed the F-actin bundling activity of HYC2 (Figure 4E upper panel). After co-incubation of F-actin and HYC2 *in vitro*, bundled F-actin filaments were observed in the presence of GST- and 6XHis-tagged recombinant HYC2 but not denatured HYC2 (Figure 4E lower panel). The above results suggest that HYC2 is an F-actin binding and can bundle F-actin into higher-order structures *in vitro*.

Given that HYC2 can interact with microfilaments and microtubules, we speculated that it might interact with both simultaneously. Indeed, incubating microfilaments and microtubules with HYC2 resulted in bundle formation (Figure 4F upper panel). *In vitro* immunofluorescence analyses confirmed the co-localization of HYC2 with cross-linked microfilaments and microtubules (Figure 4F) in contrast to no primary antibody control in the lower panel of Figure 4F. All these results collectively indicate that HYC2 functions as a cytoskeletal cross-linking protein, simultaneously binding and bundling microfilaments and microtubules.

To assess the impact of HYC2’s F-actin–bundling activity on microfilament organization *in planta*, we compared F-actin organization in wild-type and *hyc2-2/-* embryos by staining F-actin with Alexa Fluor 488 phalloidin. Wild-type globular embryos exhibited strong and elongated fluorescent signals, whereas F-actin structures in *hyc2-2/-* embryos appeared shorter with cytoplasmic decorated punctate structures (Figure S6A). To quantify the extent of F-actin bundling in globular embryos, we measured the skewness, kurtosis, and occupancy as previously described (Figure S5B; Higaki et al., 2010). Quantitative analysis revealed reduced alignment of actin bundles in *hyc2-2* mutants, as indicated by decreased skewness and kurtosis. Additionally, the density of F-actin was higher in *hyc2-2/-* than in wild-type embryos, as evidenced by increased F-actin occupancy (Figure S6B). The distortion of F-actin organization in *hyc2-2/-* embryos underscores the role of HYC2 in F-actin bundling.

### The absence of HYC2 resulted in faulty phragmoplasts in globular embryos of Arabidopsis

In plant cell division, microtubules play a more crucial role than microfilaments, as indicated by drug treatment and molecular studies (Higaki et al., 2008; Li et al., 2015). We examined the organization of microtubules in *hyc2-2/-* embryos, particularly during cell plate formation. Compared to wild-type cells, *hyc2-2/-* cells displayed abnormal cell division without overt irregularities in cortical microtubules (Figure 5A-B). To assess the impact of HYC2 in cell division, we observed the plant-specific cytoskeletal structures known as phragmoplasts by using mGFP:AtTUA6 (Liao and Weijers, 2018) driven by an embryo-specific *WOX2* promoter in arrested globular embryos of *hyc2-2/-* compared to wild-type control embryos (Figure 5C-F).

**Figure 5.**
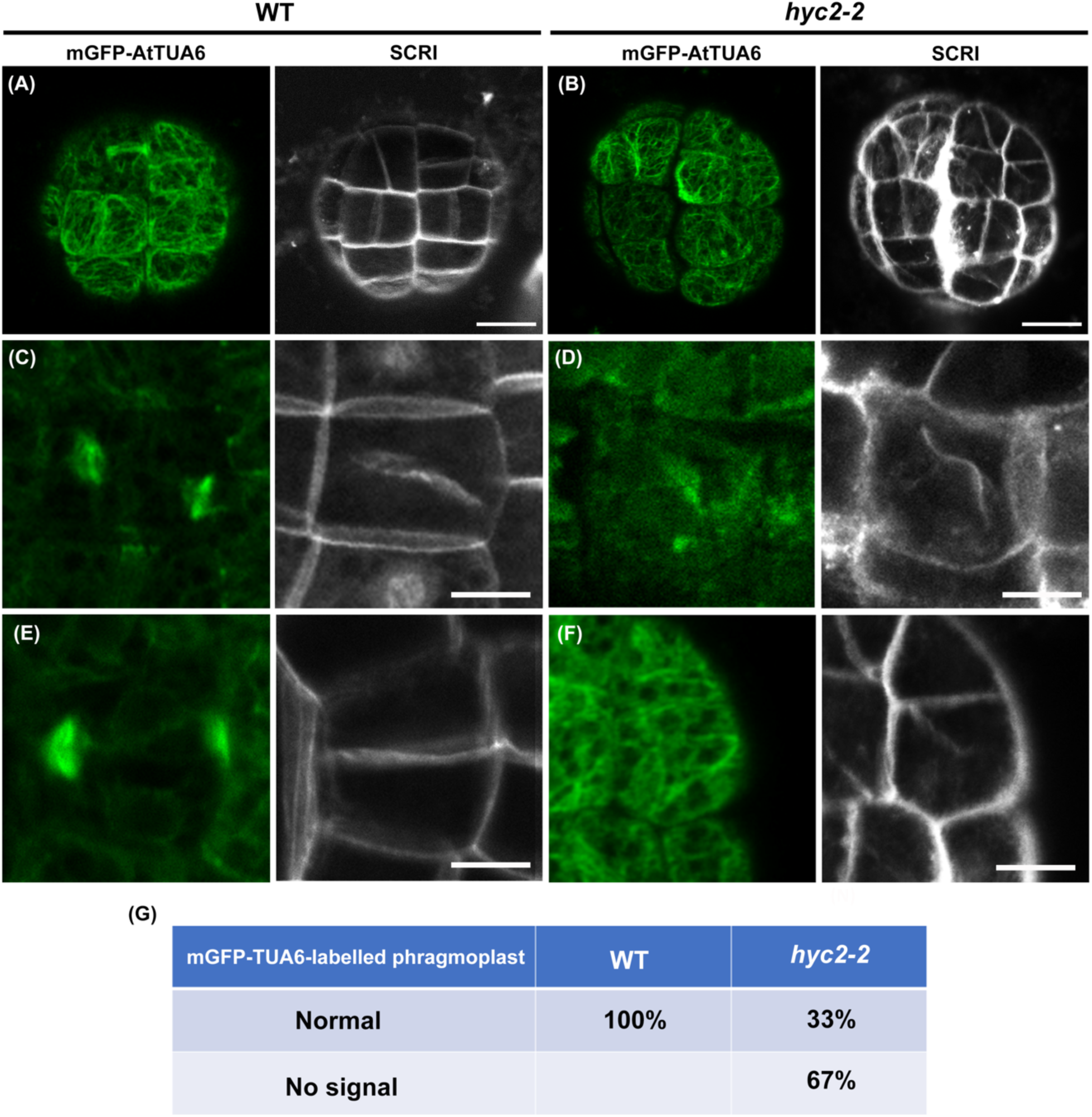
Disruption of microtubule-based phragmoplasts in *hyc2-2/-* globular embryonic cells. (A-F) Microtubules labeled with mGFP-AtTUA6 in globular embryos of the WT (A, C, E) and *hyc2-2/-* (B, D, F). Phragmoplast morphology, observed by mGFP-AtTUA6 and SCRI-labeled cell wall in embryonic cells of the wild type (C, E) and *hyc2-2/-* (D, F). Bar = 10 μm. (G) Quantitative analysis of phragmoplasts in dividing embryonic cells expressing mGFP-AtTUA6 for the WT (n=18) and *hyc2-2/-* (n=12).

At the later stages of cell plate formation, as the tubular structure of the cell plate develops, microtubule bundles typically extend toward the growing margin of the cell plate until cytokinesis is completed (Figure 5C, E). We identified dividing cells through SCRI-stained floated cell plates (Figure 5C, D) and cell wall stubs in *hyc2-2/-* cells (Figure 5F). In wild-type dividing cells, the microtubule-based phragmoplast accumulated at the leading edge of the cell plate (Figure 5C). However, in approximately 67% of dividing cells in *hyc2-2/-*, the microtubule signals of the phragmoplast were absent (Figure 5G). In all *hyc2-2/-* embryonic cells with cell wall stubs, no phragmoplast signals labeled by mGFP-TUA6 were detected (Figure 5F). Our findings suggest that HYC2 is essential for microtubule organization during cell plate formation. HYC2, with its cytoskeleton-bundling function, regulates microtubule organization in cell plate formation.

### HYC2 interacts with SH3P2 and DRP1A and is crucial for recruiting SH3P2 and DRP1A to the cell plate

To understand the role of HYC2 in cell plate formation, we investigated its protein– protein interactions with other known participants in cell plate formation by using yeast two-hybrid (Y2H) methods (Figure 6A, Figure S7). HYC2 exhibited a weak interaction with SH3P2, which is involved in membrane tubulation (Figure 6A). The HYC2–SH3P2 interaction was further confirmed by co-IP (Figure 6B). Because SH3P2, forming a complex with DRP1A, plays a vital role in establishing a tubular-vesicular network (Ahn et al., 2017), we also explored the interaction between HYC2 and DRP1A. DRP1A did not interact with HYC2 in the Y2H system, whereas co-IP assays (Figure 6B) and co-IP/LCMSMS analysis (Table S3) revealed an interaction between HYC2 and DRP1A. Thus, HYC2 likely forms a complex with SH3P2 and DRP1A during cell plate formation.

**Figure 6.**
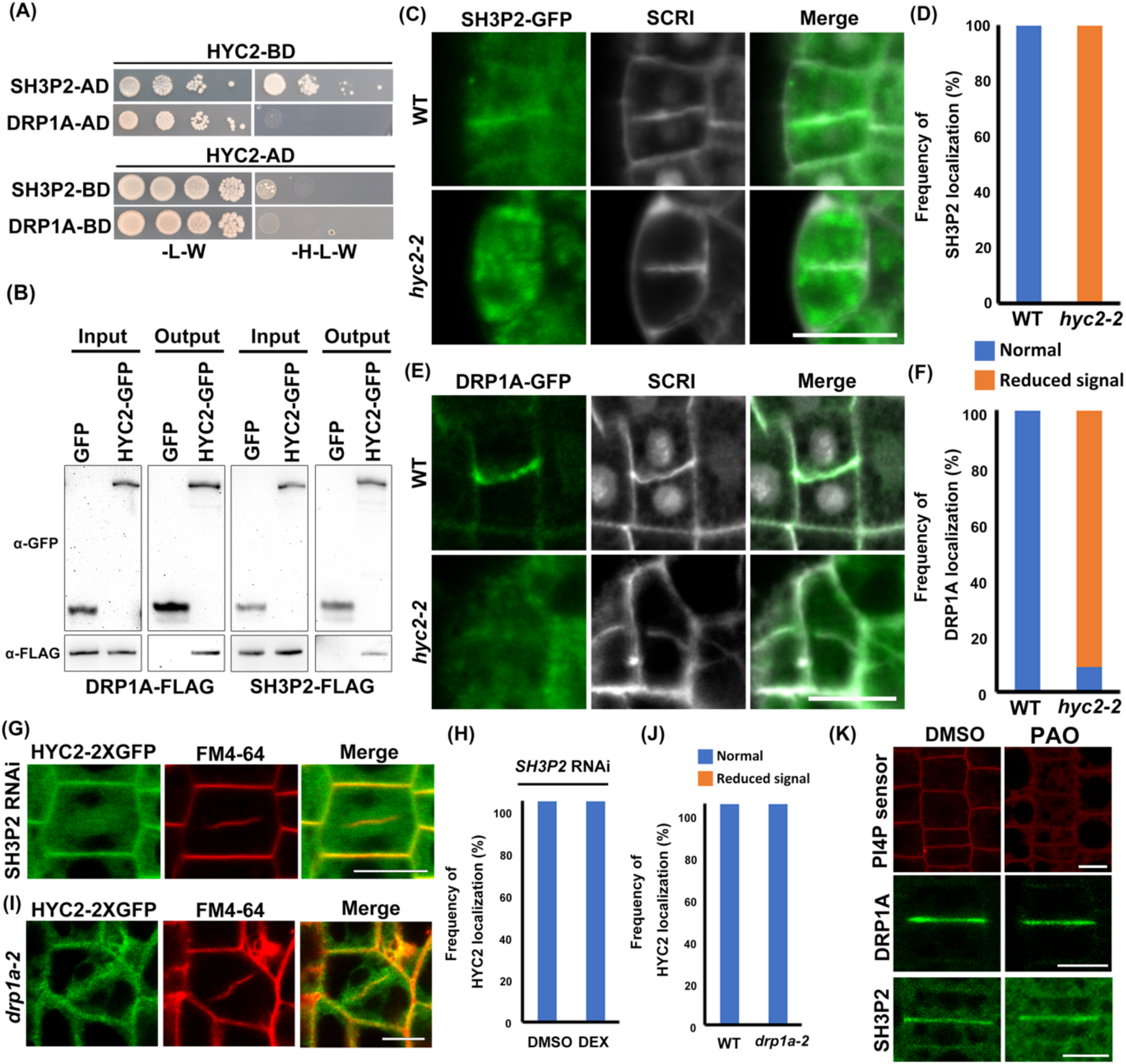
HYC2 plays an essential role in recruiting SH3P2 and DRP1A for the development of cell plates. (A) A Yeast two-hybrid assay examining protein interactions between HYC2 and SH3P2/DRP1A. Growth on synthetic dropout (SD) medium lacking tryptophan (W) and leucine (L) indicates yeast cell viability, while additional histidine (H) depletion monitors the tested protein’s interactions. (B) Co-immunoprecipitation analysis of HYC2 and SH3P2/DRP1A protein interactions. Protein extracts (Input) from transiently expressed HYC2-GFP and DRP1A/SH3P2-FLAG in Arabidopsis leaf protoplasts were subjected to GFP TrapA beads. Precipitated proteins (Output) were immunoblotted with anti-GFP and anti-FLAG antibodies. (C) Accumulation of SH3P2-GFP in the cell plate of WT and hyc2-2 globular embryos. (D) Quantitative analysis of SH3P2 signals in cell plates of WT and hyc2-2 embryos expressing SH3P2-GFP (n=19 for the WT and n=19 for hyc2-2). (E) Accumulation of DRP1A-GFP in the cell plate of WT and *hyc2-2/-* globular embryos. Cell walls were stained with SCRI. (F) Quantitative analysis of DRP1A signals in cell plates of WT and *hyc2-2/-* embryos expressing DRP1A-GFP (n=19 for the WT and n=36 for *hyc2-2/-*). (G) Localization of HYC2-2XGFP in dividing root cells of 7-day-old *SH3P2* RNAi seedlings under 10 μM dexamethasone treatment. The plasma membrane and dividing cell plate were labeled with FM4-64. Bar = 10 μm. (H) Quantitative analysis of HYC2-2XGFP signals in the cell plate of *SH3P2* RNAi seedlings with or without dexamethasone (DEX) treatment (n=27 for DMSO and n=29 for DEX). (I) Localization of HYC2-2XGFP in dividing root cells of 7-day-old *drp1a-2* mutants. The cell membrane and dividing cell plate were labeled with FM4-64. Bar = 10 μm. (J) Quantitative analysis of HYC2-2XGFP signals in the cell plate of *drp1a-2* seedlings (n=16 for the WT and n=18 for *drp1a-2*). (K) Localization of the PI4P sensor (PH domain of FAPP1), DRP1A-GFP (n=17 for DMSO and n=30 for PAO treatment), and SH3P2-GFP (n=15 for DMSO and n=15 for PAO treatment) in dividing root cells under treatment with 10 μM PAO or DMSO for 30 min. Bar = 10 μm.

To investigate the impact of HYC2 on SH3P2 and DRP1A localization, we generated DRP1A-GFP and SH3P2-GFP transgenic plants and observed their signals in the dividing cells of *hyc2-2/-* globular embryos. SH3P2 signals driven by its native promoter accumulated in the cell plate in wild-type embryos but were reduced in *hyc2-2/-* globular embryos (Figure 6C-D). Similarly, DRP1A-GFP signals in wild-type dividing cells were substantially accumulated in the cell plate (Figure 6E), whereas approximately 89% of dividing cells in *hyc2-2/-* globular embryos showed reduced DRP1A signals (Figure 6F). In *hyc2-2/-* cells, SH3P2 and DRP1A signals were dispersed in the cytosol and occasionally accumulated in dot-like structures (Figure 6C, E). This finding suggests that the cytoskeleton-associated HYC2 may contribute to the localization of SH3P2 and DRP1A to the cell plate, and the leaky cell plate phenotype in *hyc2-2/-* may result from reduced signals of SH3P2 and DRP1A, thus preventing the formation of a stable tubulo-vesicular network structure.

Conversely, we explored whether HYC2 localization depended on DRP1A and SH3P2 during cell plate formation. *SH3P2* RNAi and *drp1a-2* T-DNA mutants have defects in cell plate formation, including irregular cell organization and cell wall stubs found in *hyc2-2/-* globular embryos (Mravec et al., 2011; Ahn et al., 2017). Loss of SH3P2 and DRP1A in *SH3P2* RNAi plants and *drp1a-2* T-DNA mutants did not affect HYC2 localization (Figure 6G-J), which suggests that HYC2 localization may be independent of DRP1A and SH3P2. HYC2’s role in regulating phragmoplast formation and its impact on SH3P2 and DRP1A localization to the cell plate underscores its vital role in cell plate formation. The phytohormone auxin plays a critical role in establishing embryo patterns, regulating cell division, and determining cell identity in plants (Blilou et al., 2005; Moller and Weijers, 2009; Schlereth et al., 2010; Lau et al., 2012). PIN-FORMED (PIN) proteins, particularly PIN1, contribute to establishing auxin gradients in globular embryos. PIN1 and DRP1A form transient complexes during cell plate formation, and the polarization of PIN1 depends on DRP1A (Mravec et al., 2011). Observing that *hyc2-2/-* embryos failed to progress to the heart stage, we examined auxin gradients and PIN1 localization in globular embryos by crossing a *DR5* reporter and PIN1-GFP with *hyc2-2/+* mutants. In the wild type, the *DR5* reporter was expressed in the apical cell and its descendants until the 32-cell stage, after which *DR5* activity shifted to the uppermost suspensor cells (Friml et al., 2003). Although *DR5* activity was present in the uppermost suspensor cells in *hyc2-2* globular embryos, auxin signals abnormally accumulated in the upper tier and part of the lower tier (Figure 7), thus indicating altered auxin distribution in the *hyc2-2/-* mutant. In the wild-type globular embryo, PIN1 signals accumulated at the plasma membrane of the upper tier and provascular cells. PIN1-GFP still localized to the plasma membrane in *hyc2-2/-* mutants but exhibited a slight increase in cytosolic accumulation (Figure 7). These findings suggest that the developmental defects in *hyc2-2/-* embryos are linked to abnormal auxin distribution and PIN1 polarization. Because PIN1 recruitment to the plasma membrane remains intact, HYC2 is critical for recruiting SH3P2 and DRP1A to the cell plate instead of PIN1 trafficking along the secretory pathway.

**Figure 7.**
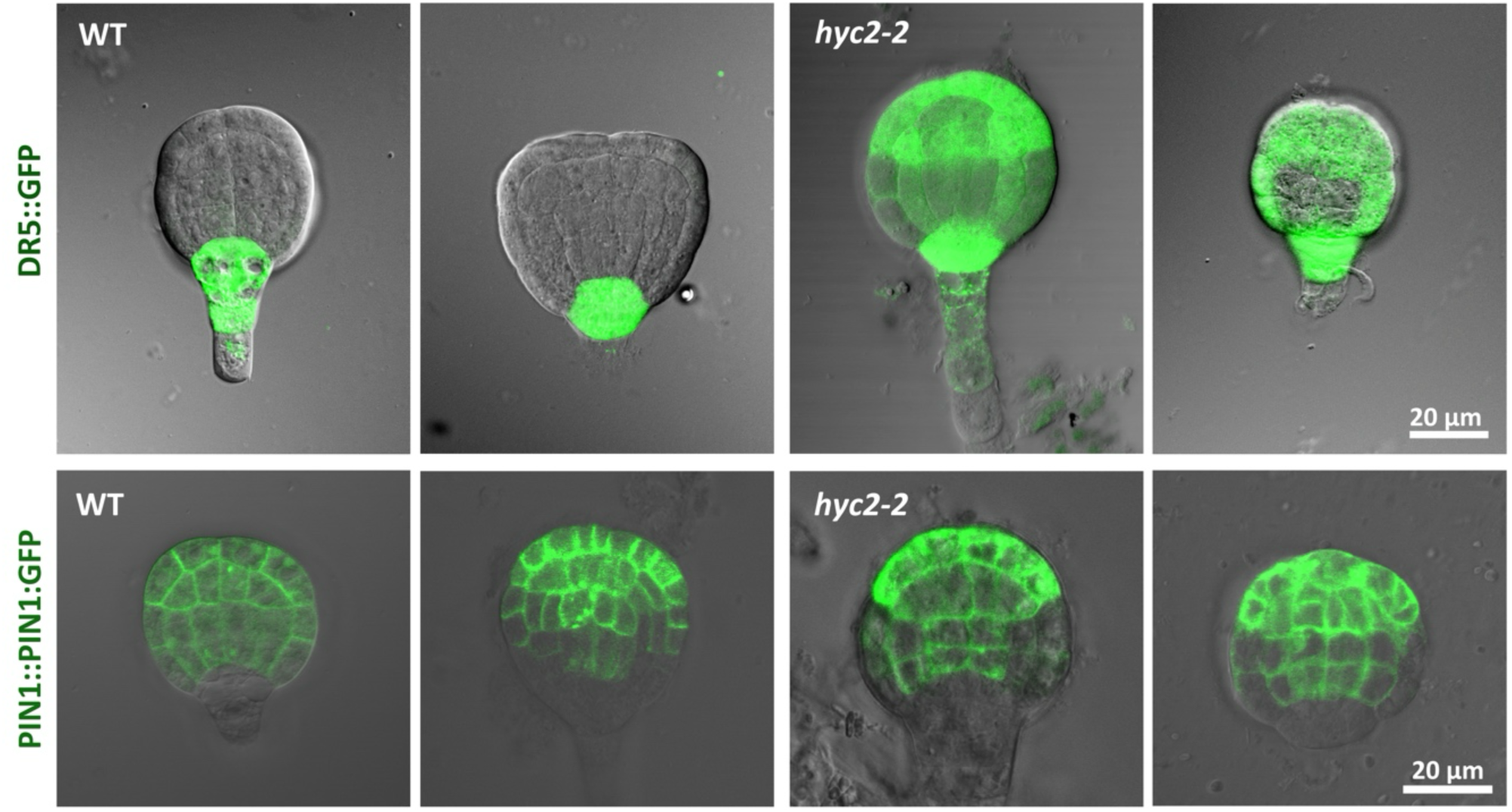
The disruption of the auxin gradient and the alteration in auxin transport polarity in *hyc2-2/-* mutants. Observation of auxin distribution and PIN1-GFP expression in Arabidopsis embryos with *DR5::GFP* (n=12 for the WT and n=18 for *hyc2-2/-*) and PIN1-GFP (n=7 for the WT and n=20 for *hyc2-2/-*) signals by confocal microscopy. Bar = 20 μm.

## Discussion

We elucidated the molecular role of HYC2 as a cytoskeleton-bundling protein crucial for cell plate formation during early Arabidopsis embryogenesis. The HYC2 mutation resulted in cytokinesis defects, forming arrested globular embryos and aborted seeds. After metaphase, HYC2 exhibited accumulation around the FM4-64-labeled cell plate and phragmoplasts. Depletion of HYC2 after anaphase caused embryo arrest, which highlights the indispensable role of HYC2 in cell plate formation. Moreover, HYC2 played a regulatory role in phragmoplast formation and influenced the recruitment of SH3P2 and DRP1A to the cell plate. These findings underscore the essential role of HYC2, with its cytoskeleton-bundling activity, in phragmoplast formation and the recruitment of SH3P2 and DRP1A to the developing cell plate.

FAM126A, a human protein analogous to HYC2, interacts with the PI4K complex to regulate PI4P production in humans. However, the precise mechanism governing the association between FAM126A and the PI4K complex remains unclear (Yao et al., 1999; Stevenson-Paulik, Love and Boss, 2003). A recent study indicated that HYC2 is a constituent of the PI4Kα1 complex and localizes to Arabidopsis plasma membrane nanodomains (Noack et al., 2022). Notably, except for *hyc2* mutants, mutants of various other components, including PI4Kα1, HYC1, NPG proteins, and EFOP (EFR3-OF-PLANTS) proteins, exhibit gametophyte-defective phenotypes (Noack et al., 2022). Loss of HYC2 leads to embryo lethality at the globular stage in Arabidopsis. These lethal phenotypes pose challenges in exploring the function of the PI4Kα1 complex in plants. Furthermore, the molecular mechanism and biological role of HYC2 remain unexplored, even in animals. Our study initiates the investigation of the Hyccin family, shedding light on its role in regulating cytoskeleton dynamics during cell plate formation and influencing the recruitment of proteins to the cell plate in Arabidopsis.

Cell plate formation relies on lipid determinants that delineate plasma membrane identity. These determinants encompass phosphatidylinositol 4-phosphate (PI4P), synthesized by PI 4-kinases, including PI4Kα1, PI4Kβ1, and PI4Kβ2 (Hammond et al., 2012; Nakatsu et al., 2012; Baskin et al., 2016). PI4Kα1 primarily functions at the plasma membrane, whereas PI4Kβ1 and PI4Kβ2 generate the PI4P pool at the *trans*-Golgi network (Preuss et al., 2006; Lin et al., 2019). PI4Kβ1 and PI4Kβ2 play roles in cell plate formation, as evidenced by the *pi4kβ1pi4kβ2* mutant displaying over-stabilized phragmoplast microtubules, mis-localization of MAP65-3 and defects in the membrane trafficking of KNOLLE and PIN2 (Lin et al., 2019). In contrast, our study revealed that HYC2, a component of the PI4Kα1 complex, protects microtubules against disassembly, and *hyc2-2/-* embryonic cells lose microtubule-based phragmoplasts. Despite PI4Kβ1/PI4Kβ2 and HYC2 influencing membrane trafficking, they appear to regulate phragmoplasts by distinct mechanisms. In yeast, PI4Kα and PI4Kβ groups are non-redundant, with no substitution possible between them (Audhya et al., 2000). Our findings support that these two groups are not functionally redundant in plants.

The creation of the cell plate encompasses the incorporation of membrane structures into an existing plasma membrane, a process tightly governed by the secretory system’s regulation of materials delivered to the developing cell plate (Backues et al., 2007). This regulation is intricately linked to the organization of microtubules in phragmoplasts and microtubule-associated proteins, including the Kinesin-12 subfamily and MAP65-3, which are associated with the phragmoplast. Kinesin-12 proteins are recognized for their role in transporting cargo along microtubules (Muller and Livanos, 2019). KNOLLE, a syntaxin-like protein, is implicated in cytokinesis (Lauber et al., 1997), and in *kin12akin12b* double mutants, the failure to organize microtubule bundles into an antiparallel phragmoplast array resulted in the loss of KNOLLE’s localization to the cell plate (Lee, Li and Liu, 2007). Conversely, MAP65-3 influenced microtubule-plus ends in phragmoplasts but did not affect the membrane trafficking of KNOLLE and DRP1A (Ho et al., 2011). These observations suggest that cytoskeleton-associated proteins play crucial roles in phragmoplast organization but may not be directly involved in transporting cargo to the cell plate. Notably, our results demonstrate that cytoskeleton-associated HYC2 regulates both the formation of phragmoplasts and the delivery of cargo to the cell plate during cell plate formation.

To figure out if the recruitment is related to PI4P production, we treated Arabidopsis seedlings of DRP1A-GFP transgenic plants with phenylarsine oxide (PAO), PI4P inhibitor to detach PI4P from plasma membrane to cytosol. Our results showed that both DRP1A and SH3P2 signals were still in the cell plate after PAO treatment in contrast to the PI4P sensor, mRFP-PH_FAPP1_ signals were released from the plasma membrane to the cytosol (Figure 6K). These suggest that PI4P signals have less impact on targeting DRP1A and SH3P2 to the cell plate. However, we could not rule out the effect of PI4P on DRP1A and SH3P2 because both DRP1A-GFP and SH3P2-GFP fluorescent intensity were slightly reduced after PAO treatment, and previous studies showed that SH3P2 binds to PI4P (Zhuang et al., 2013; Mravec et al., 2011; Ahn et al., 2017). Accordingly, the cytoskeleton binding function of HYC2 plays a critical role in SH3P2 and DRP1A recruitments during cell plate formation.

Using biochemical interaction studies and *in planta* investigations, we observed impaired localization of SH3P2 and DRP1A in incomplete cell plates in *hyc2-2/-* embryonic cells. These findings indicate that HYC2, with its cytoskeleton-bundling activity, is essential for adequately localizing SH3P2 and DRP1A to the cell plate. Previous research has demonstrated the interaction between DRP1A and PIN1 in correcting the PIN1 distribution (Mravec et al., 2011). Notably, in *hyc2-2/-* mutants, we did not observe a depletion of PIN1 at the plasma membrane, although DRP1A localization was altered. HYC2 plays a role in both phragmoplast formation and the trafficking of cargo, including SH3P2 and DRP1A, to the cell plate. Given that HYC2 is a part of the PI4Kα1 complex, further investigation into the relation between the PI4Kα1 complex and the SH3P2–DRP1A complex promises to be intriguing.

## Methods

### Plant material and growth conditions

All Arabidopsis plants and materials used in this study were in the Col-0 ecotype background. T-DNA insertion mutants of *hyc2-2* (SALK_003807) and *hyc2-3* (SALK_040977) were obtained from the Arabidopsis Biological Resource Center (Ohio State University, USA). Seeds from *Arabidopsis thaliana* were surface-sterilized after stratification for 3 days at 4°C and germinated on half-strength Murashige and Skoog (MS) medium (Duchefa Biocheme, The Netherlands) containing 1% sucrose and 0.7% phytoagar (Duchefa Biocheme, The Netherlands) or grown in the soil inside a growth chamber with a 16-h light/8-h dark cycle at 22°C.

### Confocal microscopy and imagining

Microscopic analysis of embryonic cells from *hyc2-2* or transgenic plants carrying SH3P2-GFP, DRP1A-GFP, *DR5*::GFP, PIN1-GFP, and mGFP:AtTUA in *hyc2-2* were observed under a Zeiss LSM880 confocal microscope. The outline of the embryonic cell wall was visualized by SCRI staining. Primary roots from transgenic plants with HYC2-2XGFP were stained with FM4-64 (Invitrogen) for root cells. GFP signals from HYC2, FM4-64/PI, and SCRI were recorded using GFP, rhodamine, and DAPI filters. Line graphs of the signal intensities were processed by using ImageJ.

### Ultrastructural analysis

For transmission electron microscopy (TEM), developing embryos of Arabidopsis were dissected from siliques. The samples were fixed in 2.5% glutaraldehyde and 4% paraformaldehyde in 0.1 M sodium phosphate buffer, pH 7.0, at room temperature for 4 h. After three 20-min buffer rinses; the samples were post-fixed in 1% OsO_4_ in the buffer for 4 h at room temperature and rinsed for 20 min three times. The samples were then dehydrated in an acetone series, embedded in Spurr’s resin, and sectioned with a Lecia Reichert Ultracut S or Lecia EM UC6 ultramicrotome. The ultra-thin sections (70-90 nm) were stained with uranyl acetate and lead citrate. A Philips CM 100 transmission electron microscope at 80 KV was used for viewing, and images were captured using a Gatan Orius CCD camera.

### Globular embryo observation

To examine the early embryogenesis of *hyc2* and *HYC2pro::HYC2-CYCB1;2-2XGFP*/*hyc2-2/+* transgenic plants, young siliques were fixed in ethanol: acetic acid (8:1) for 30 min. They were cleared in Hoyer’s solution (chloral hydrate: distilled water: glycerol=8:3:1). Developing seeds were dissected from siliques and photographed by using an Olympus BX51 camera and a Zeiss Axio Imager Z1 Microscope.

### Co-immunoprecipitation, LC-MS/MS analysis, and immunoblot analysis

Cell lysates were obtained from 5 g of 7-day-old seedlings harboring *HYC2p:HYC2-2XGFP* and *GFP* in extraction buffer (100 mM Tris, pH 8.0, 150 mM NaCl, 5 mM EDTA, 5 mM EGTA, 10 mM DTT, 0.1% NP-40, and protease inhibitor cocktail [Roche]). Then, 50 μl GFP-Trap A (Chromo-Tek) was washed with extraction buffer and added to cell lysates for incubation at 4°C for 2 h with gentle rotation. After washing three times with extraction buffer, proteins were eluted for LC-MS/MS analysis to identify the potential interacting proteins. The acquired MS/MS data were analyzed using the SEQUEST search program (BioWorks Browser 3.3, Thermo Fisher Scientific) against the *Arabidopsis* database downloaded from the National Center for Biotechnology Information (NCBI). Immunoblots were detected by chemiluminescence (SuperSignal West Pico, Thermo) for immunoblotting.

### Purification of HYC2 recombinant protein from E. coli

To construct the HYC2 recombinant protein, PCR products using primer sets (HYC2 ORF w/o SP BamHI F and HYC2 ORF SalI R; HYC2 ORF-F BamH1 and HYC2 ORF-R XhoI) were ligated into the vector pGEX 4T-1 and pET30a to generate a fusion protein carrying the GST and 6XHis-tag, respectively. The recombinant proteins were expressed in *E. coli* strain BL21 and incubated with resins (GE Healthcare) according to the manufacturer’s instructions. The resins were intensively washed with incubation buffer before eluting the recombinant proteins. The eluted protein was further buffer-exchanged using a PD-10 column (GE Healthcare).

### Microscale thermophoresis (MST) measurements

MST analysis was performed using a Microscale Thermophoresis Monolith NT.115Pico instrument. Actin, Tubulin, or HYC2 was labeled with Alexa-647 using a Monolith NT™ Protein Labelling Kit RED-NHS. The protein interaction was determined after 10 min incubation at room temperature in Tris buffer (50 mM Tris, pH 8.0, 100 mM NaCl). Finally, 3 to 5 μl was loaded into standard capillaries, and thermophoresis was determined.

### MT binding and bundling assay

According to the manufacturer’s manual, microtubules were obtained from tubulin (T240, Cytoskeleton). All proteins were centrifuged at 100,000 × g for 30 min at 4°C before use. The reaction was buffered with 80mM PIPES, pH6.9, 2mM MgCl_2,_ 0.5mM EGTA, 10% glycerol and 1mM GTP. In the high-speed experiments, samples were centrifuged at 100,000 × g for 30 min to pellet the MTs. The presence of HYC2 in the supernatants and pellets was analyzed via SDS-PAGE and Coomassie Brilliant Blue R staining. In the low-speed experiments, the samples were centrifuged at 12,500 × g for 30 min to pellet high-order MTs. MT in the supernatants and pellets was revealed via SDS-PAGE and Coomassie Brilliant Blue R staining.

### Actin binding and bundling assay

High- and low-speed co-sedimentation experiments were performed to assess the F-actin binding and bundling activities of HYC2. All proteins were centrifuged at 100,000 × g for 30 min at 4°C before use. The reaction medium was buffered with 5 mM MES (pH 6.2) and supplemented with 10μM CaCl_2_. In the high-speed experiments, samples were centrifuged at 100,000 × g for 30 min to pellet the actin filaments. The presence of HYC2 in the supernatants (F-actin unbound fraction) and pellets (F-actin bound fraction) was analyzed via SDS-PAGE and Coomassie Brilliant Blue R staining. In the low-speed experiments, the samples were centrifuged at 12,500 × g for 30 min to pellet high-order F-actin structures. The presence of F-actin in the supernatants and pellets was analyzed via SDS-PAGE and Coomassie Brilliant Blue R staining.

### In vitro visualization of microtubules and microfilaments

The bundles of microtubules and actin filaments in the samples were also visualized directly by using a Zeiss LSM510 confocal microscope. Rhodamine-red–labeled microtubules were incubated with 1 μM HYC2-6XHis at 22°C for 30 min or on ice for 30 min. For F-actin bundles, preformed F-actin was incubated with 1 μM GST-HYC2, HYC2-6XHis, and denatured GST-HYC2 at 22°C for 30 min. The HYC2 recombinant protein was boiled for 10 min to denature the proteins. Actin filaments were stained with 350 nM Alexa 488-phalloidin (Invitrogen) and observed under the same excitation and emission conditions as GFP fluorescence under the confocal microscope.

### In vitro immunofluorescence

*In vitro* immunofluorescence was performed to investigate the location of HYC2 with cytoskeleton bundles. Preformed microfilaments and rhodamine-red–labeled microtubules were incubated with HYC2-6XHis at room temperature for 5 min. Anti-6XHis antibody (Invitrogen 46-0693, 1:50) and anti-mouse IgG-TRITC antibody (Sigma T5393, 1:50) were added by incubation at room temperature for 15 min. Actin filaments were stained with 350 nM Alexa 633-phalloidin (Invitrogen). HYC2 incubated with a secondary antibody alone was a negative control. Aliquots were transferred to slides pretreated with poly-L-Lys and visualized by confocal microscopy.

### Antibodies

The sources for the antibodies are as follows, with the working dilution in parentheses: GFP (1:1000, Santa Cruz, sc-9996), actin (1:500, Pierce, MA1-744), mouse IgG (1:1000, Jackson ImmunoResearch, 115-035-003), 6XHis (1:50, Invitrogen, 46-0693), FLAG (1:1000, Sigma, F7425) and mouse IgG-TRITC (1:50, Sigma, T5393)

## Acknowledgments

We thank Dr. Bo Liu for the helpful discussion about our study. We thank Mrs. Mei-Jane Fang (DNA Analysis Core Laboratory, Academia Sinica) for assistance with the confocal microscopy analyses and Mr. Wann-Neng Jane (Plant Cell Biology Core Laboratory, Academia Sinica) for the ultrastructural analysis. We acknowledge the use of the Microscale Thermophoresis Monolith NT.115Pico in the Biophysics Core Facility, funded by the Academia Sinica Core Facility and Innovative Instrument Project (AS-CFII-111-201). The *proDR5:GFP* and *proPIN1:PIN1-GFP* transgenic lines were generous gifts from Dr. Klaus Palme (Benkova et al., 2003). The *pWOX2: mGFP:AtTUA* transgenic lines were kindly provided by Dr. Dolf Weijers (Liao and Weijers, 2018). The construct of *pDRP1A: DRP1A* and the seeds of *drp1a-2* (SALK_06977) were kindly provided by Dr. Sebastian Y. Bednarek (Mravec et al., 2011). The seeds of *SH3P2* RNAi transgenic plants were kindly provided by Dr. Liwen Jiang (Zhuang et al., 2013). This work was supported by research grants from Academia Sinica (AS-GCS-111-L06) and the National Science and Technology Council (MOST 108-2311-B-001-036-MY and MOST 110-2313-B-001 −007 -MY3) to G.-Y. Jauh.

## Author information

Y.-W. H., H.-J.W., and G.-Y.J. conceptualized and designed the experiments. Y.-W. H., C.-L.G., and C.-T. J. conducted the experiments, and Y.-W. H. and G.-Y.J. wrote the article.

## Supplementary information

**Figure S1.**
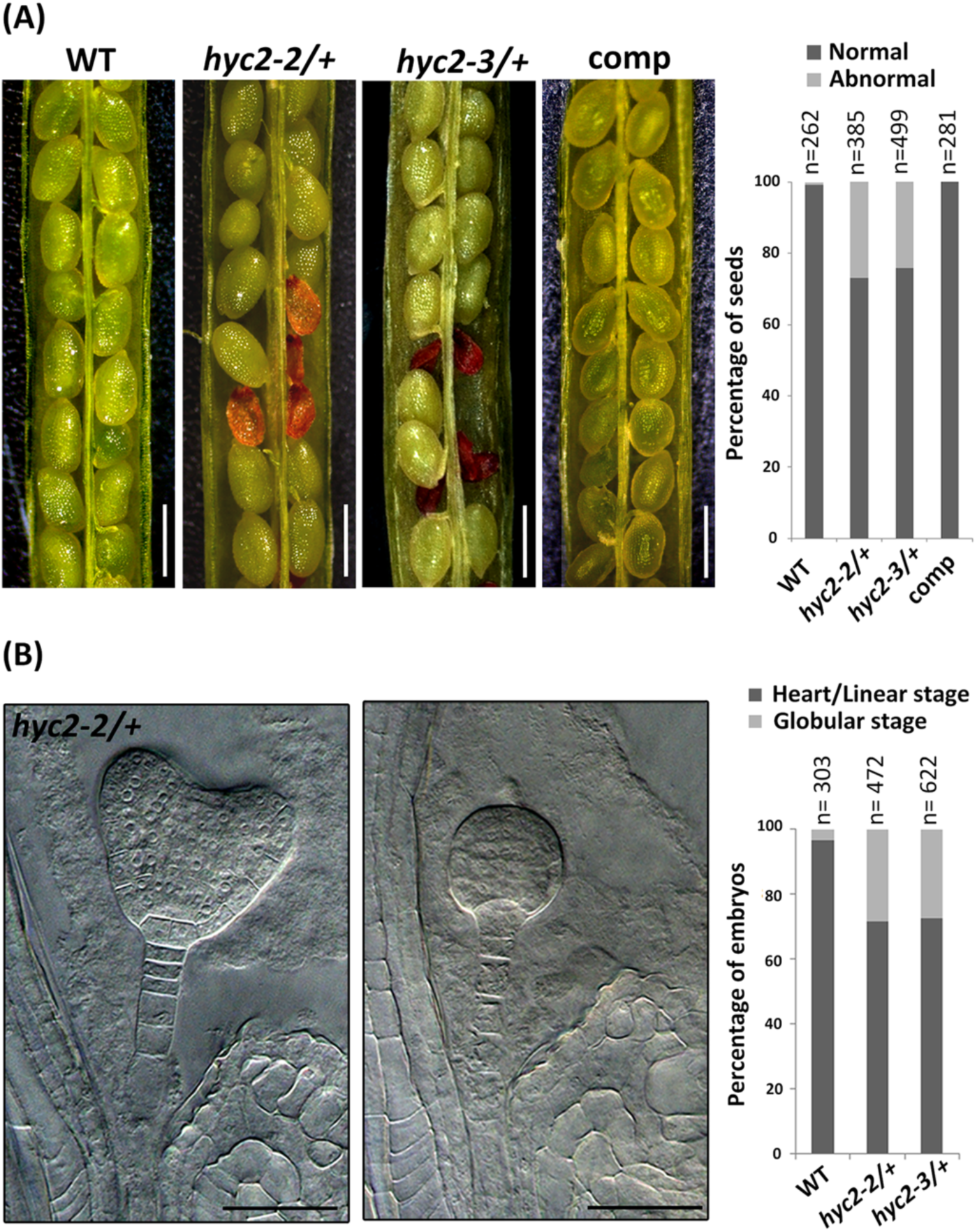
Phenotyping the *HYC2* T-DNA insertional mutants. (A) Dissected mature siliques from the wild type (WT), *hyc2-2/+*, *hyc2-3/+*, and the complementation line (comp). Plot showed the percentage of normal and abnormal seeds in dissected mature siliques, and approximately 25% of seeds from *hyc2-2/+* and *hyc2-3/+* siliques exhibit an aborted, brown, and shrunken phenotype (n=262, n=499, and n=499 for the WT, *hyc2-2/+*, and *hyc2-3/+*, respectively). *p* > 0.05 (χ2 test, WT/HZ: HM=3:1, n=3 independent experiments). Bar=500 μm. (B) Color clearance of developing siliques from *hyc2-2/+*. The plot showed the percentage of embryo developmental stage in the developing siliques, and about 25% of embryos were arrested at the globular stage. In contrast, the remaining embryos ceased development at the heart stage (n=303, n=472, and n=622 for WT, *hyc2-2/+*, and *hyc2-3/+*, respectively). *p* > 0.05 (χ2 test, WT/HZ: HM=3:1, n=3 independent experiments).

**Figure S2.**
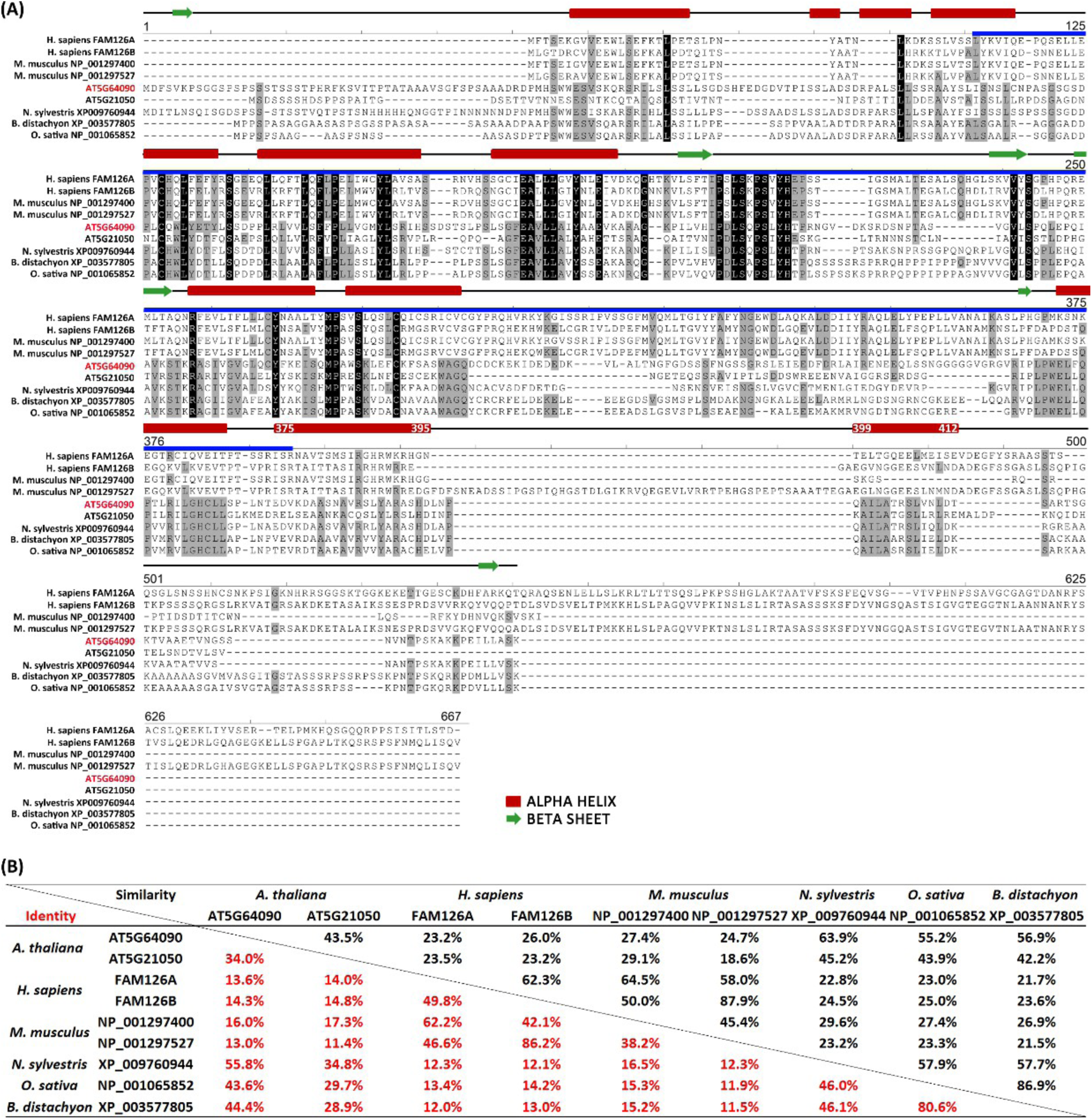
Protein characteristics of HYC2 and its homologs. (A) Alignment of protein sequences for selected Hyccin homologs from mammals and plants. AlignX generated the alignment, allowing multiple pairwise alignments to assess sequence divergence. The Hyccin domain, present in all homologs, is highlighted in blue. Identical amino acids are denoted in dark text, and conserved amino acids are shaded in grey. The secondary structure of HYC2 was predicted using JPred4 (Drozdetskiy et al., 2015) with the JNet algorithm. Alpha helices are represented by red bars, and green arrows in the JPred4 results indicate beta-helices. (B) Comparison of protein similarity and identity among HYC2 and its homologs. Percentages were determined by AlignX sequence analyses.

**Figure S3.**
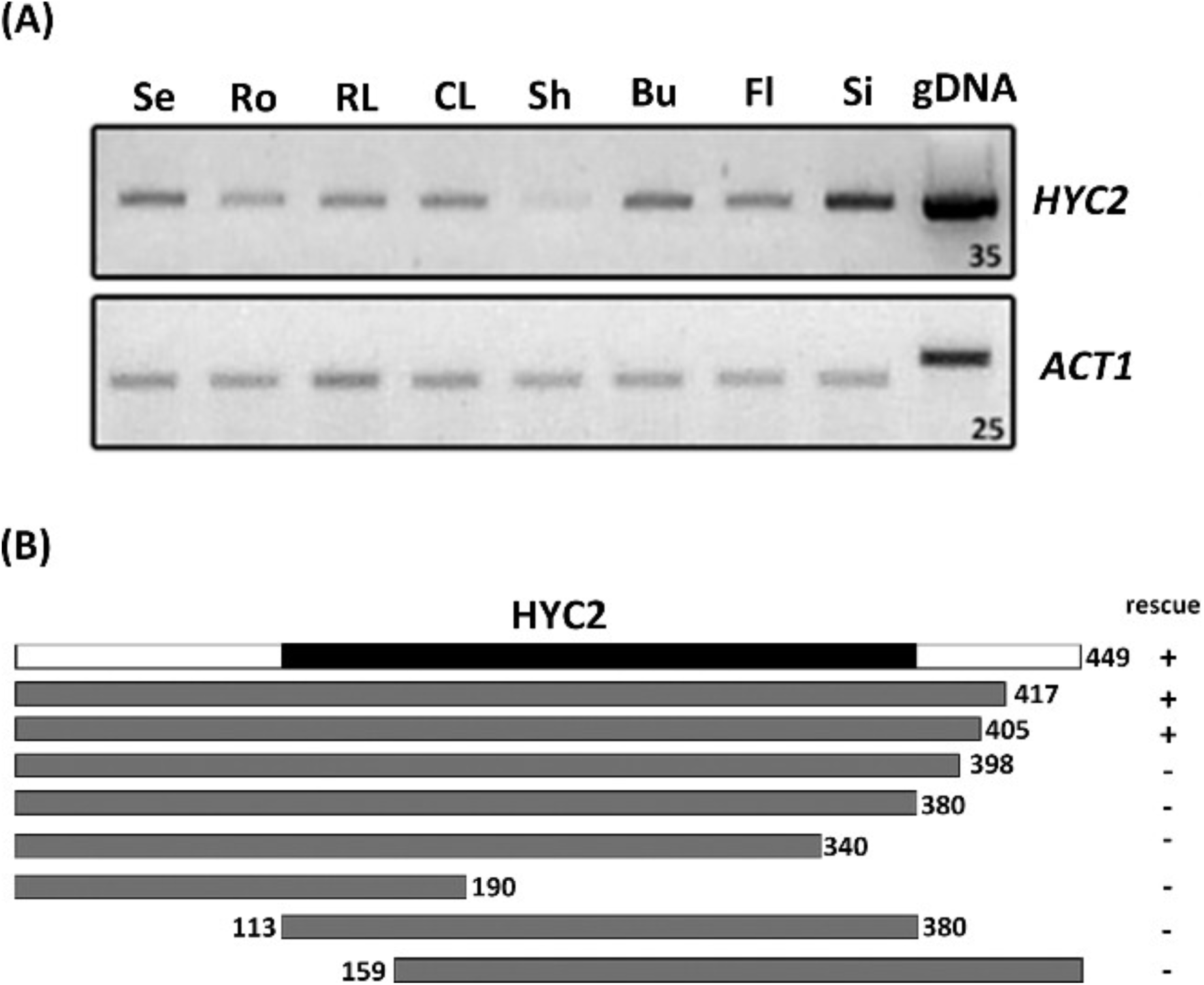
Spatial expression of HYC2 and constructs for complementation in *hyc2-2/+* mutant. (A) Quantitative PCR (qPCR) analyses of the spatial expression profile of *HYC2* in various tissues of 7-day-old seedlings (Se), roots (Ro), rosette leaves (RL), cauline leaves (CL), shoots (Sh), buds (bu), flowers (Fl), and siliques (Si), as well as genomic DNA (gDNA). *ACT1* was the internal control for normalization. The primer pairs for the internal control gene *ACT1* were designed to span introns, confirming the absence of genomic DNA contamination in all cDNA samples. The number on the right side indicates the number of amplification cycles for PCR. (B) Schematic representation of HYC2 protein variants in the complementation assay. The number of amino acids is specified, and the black region signifies the Hyccin domain.

**Figure S4.**
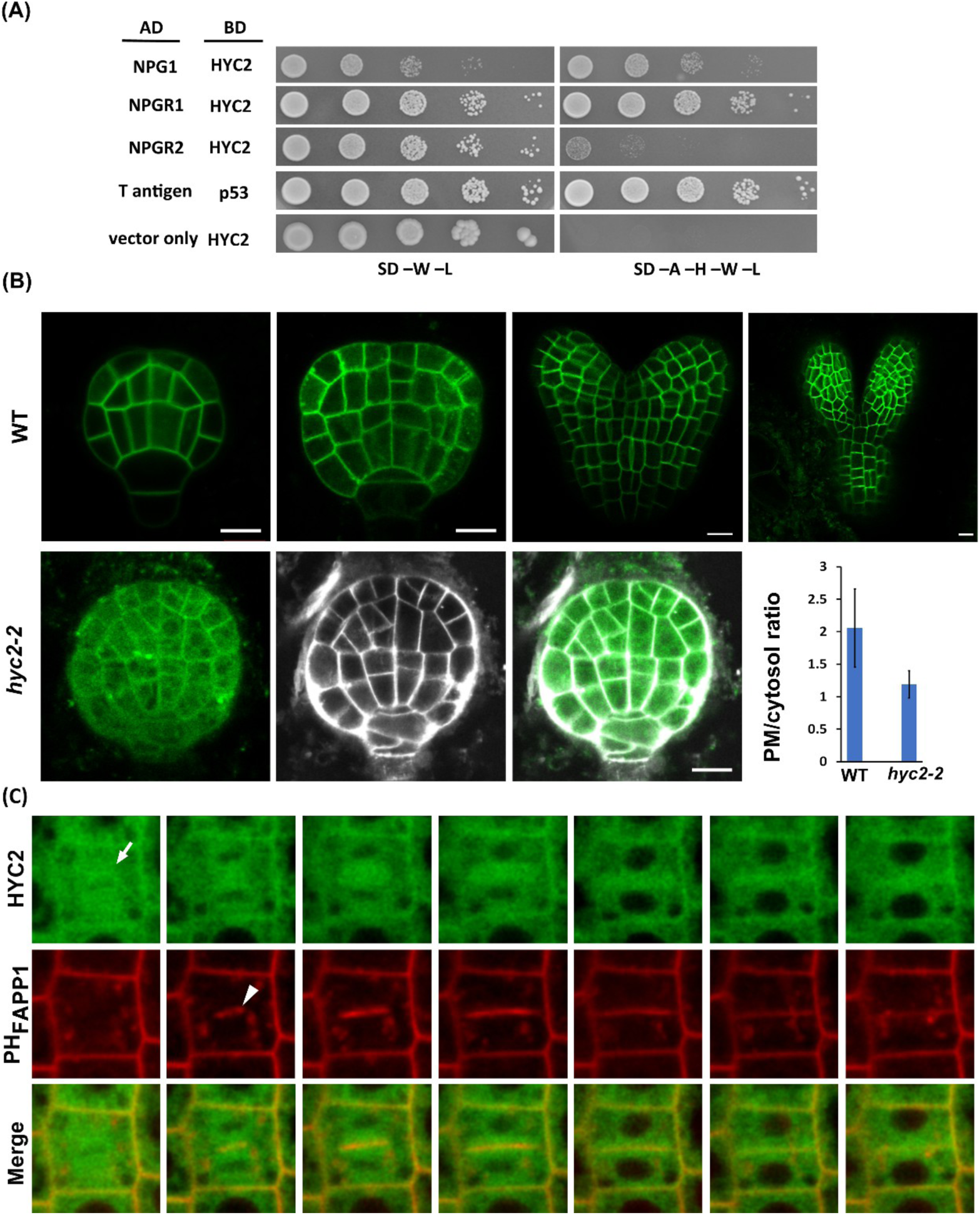
Analysis of HYC2 in PI4P signals by yeast two-hybrid (Y2H) system, HYC2-2XGFP transgenic plants, and *hyc2-2/-* mutants. (A) Investigation of protein interaction between HYC2 and NPG proteins by Y2H assay. Synthetic dropout (SD) nutrient medium lacking tryptophan (W) and leucine (L) allows yeast cells to grow regardless of protein interactions. Additional depletion of adenine (A) and histidine (H) in the SD medium was used to monitor the interaction of the tested proteins. (B) PI4P signals of YFP-PH_FAPP1_ in Arabidopsis embryos. The upper panel displays PI4P signals from wild-type (WT) embryos expressing YFP-PH_FAPP1_, and the lower panel shows PI4P signals from globular embryos of *hyc2-2/-*, along with the ratio of signal intensity from the plasma membrane to the cytosol for the PI4P biosensor YFP-PH_FAPP1_ (n=30 for WT and *hyc2-2/-*). Data are mean ± SEM. Bar = 10 μm. (C) Time-lapse imaging of root tip cells from 7-day-old seedlings carrying *HYC2-2XGFP* and the PI4P marker mRFP-PH_FAPP1_. Images were captured at 3-minute intervals. The arrow indicates HYC2 accumulation in dividing cells, and the arrowhead indicates the emergence of PI4P signals in the developing cell plate. Bar = 10 μm.

**Figure S5.**
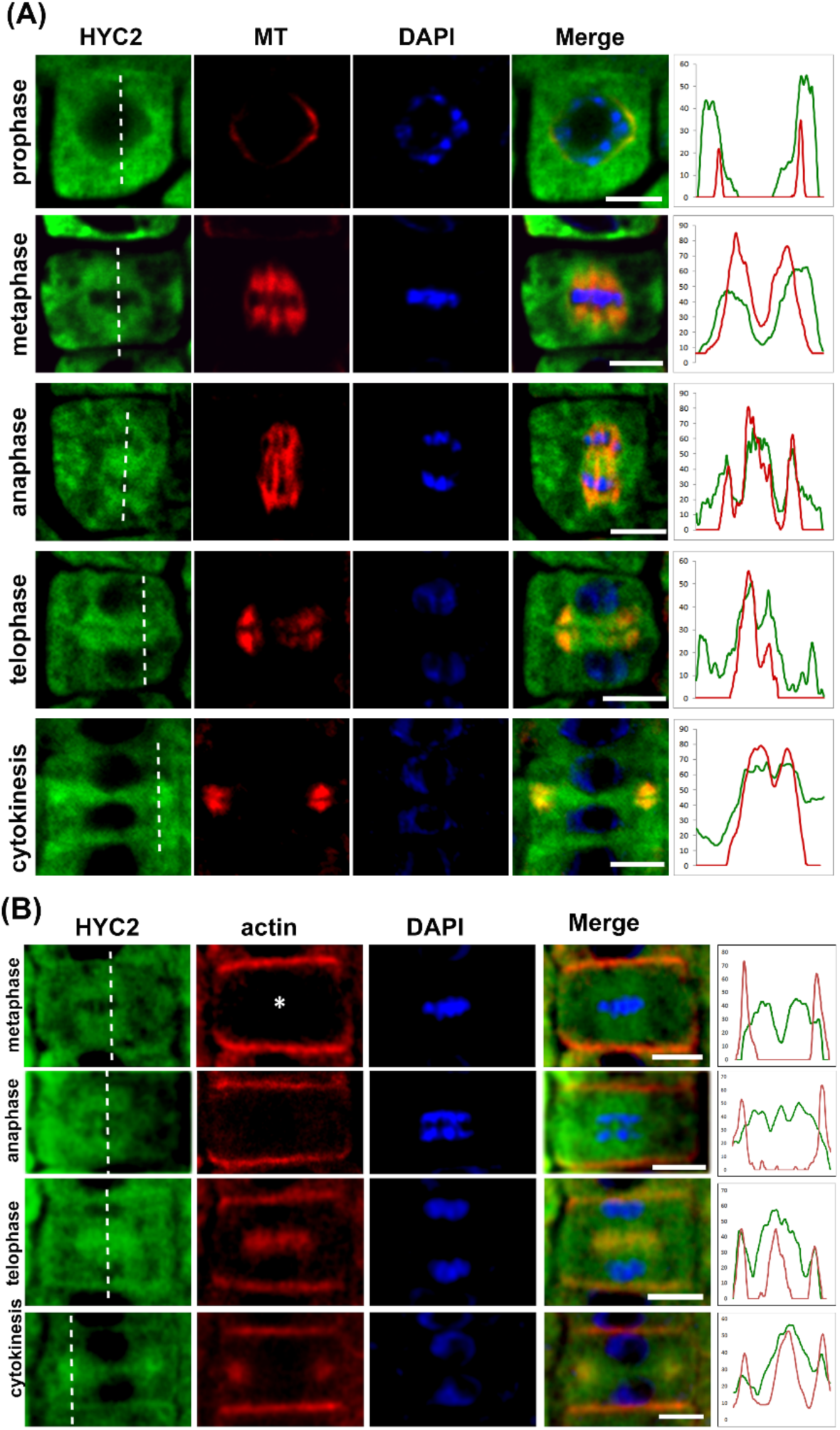
Co-localization of HYC2 with microtubules and microfilaments during mitosis. (A) Immunolocalization in root tip cells of 5-day-old seedlings with stable expression of HYC2-2XGFP using an anti-tubulin antibody (red), DAPI (blue), and anti-GFP antibody (green). Bar = 5 μm. (B) Immunolocalization in root tip cells of 5-day-old seedlings with stable expression of HYC2-2XGFP using an anti-actin antibody (red), DAPI (blue), and anti-GFP antibody (green). The line graphs in the right panels illustrate the quantified fluorescence intensity of (A) HYC2-2XGFP (green) and microtubules (red) and (B) HYC2-2XGFP (green) and F-actin (red), which correspond to the signals from the relative position indicated by the dashed line at different mitotic phases. Bar = 5 μm.

**Figure S6.**
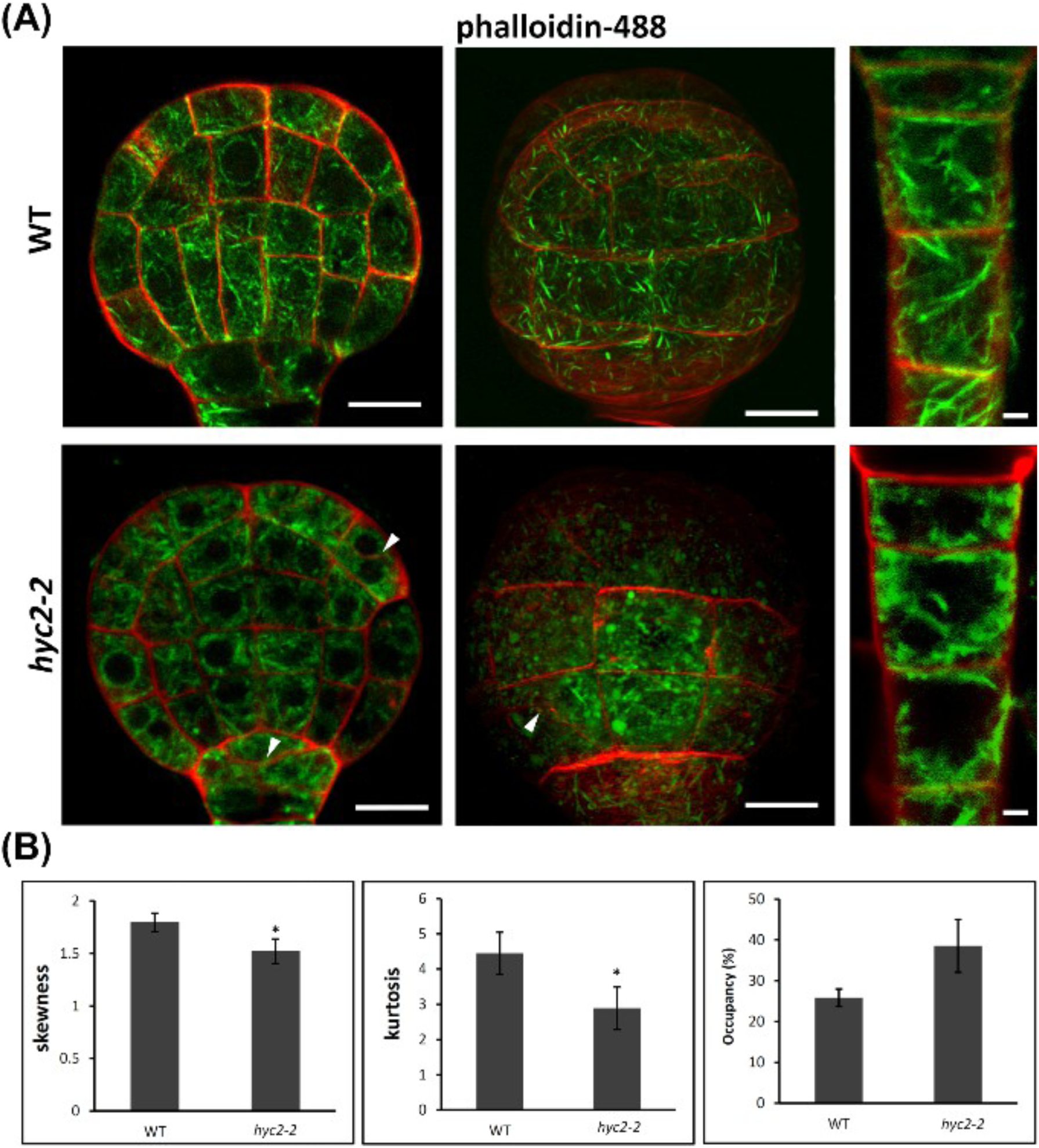
HYC2 is essential for F-actin organization in globular embryonic cells. (A) F-actin organization in globular embryos of the WT and *hyc2-2*/-. Phalloidin-stained F-actin (green) in WT and *hyc2-2/-* embryos. Red signals represent SCRI-stained polysaccharides in the cell wall, and arrowheads indicate cell wall stubs. The left panel shows longitudinal cross-sections of a globular embryo, the middle panel presents a constructed 3D image of the globular embryo, and the right panel displays a suspensor. Bar = 10 μm in the left and middle panels and 2 μm in the right panel. (B) Quantitative analysis of skewness, kurtosis, and occupancy of F-actin in WT and *hyc2-2/-* embryos. Fluorescent signals of phalloidin-stained F-actin from longitudinal cross-sections of globular embryos were captured by fluorescence microscopy. Images from globular embryos of the WT (n=21) and *hyc2-2/-* (n=20) were analyzed with ImageJ (Data are mean±SEM, *p*<0.0001, two-tailed, Student *t* test, n=3 independent experiments).

**Figure S7.**
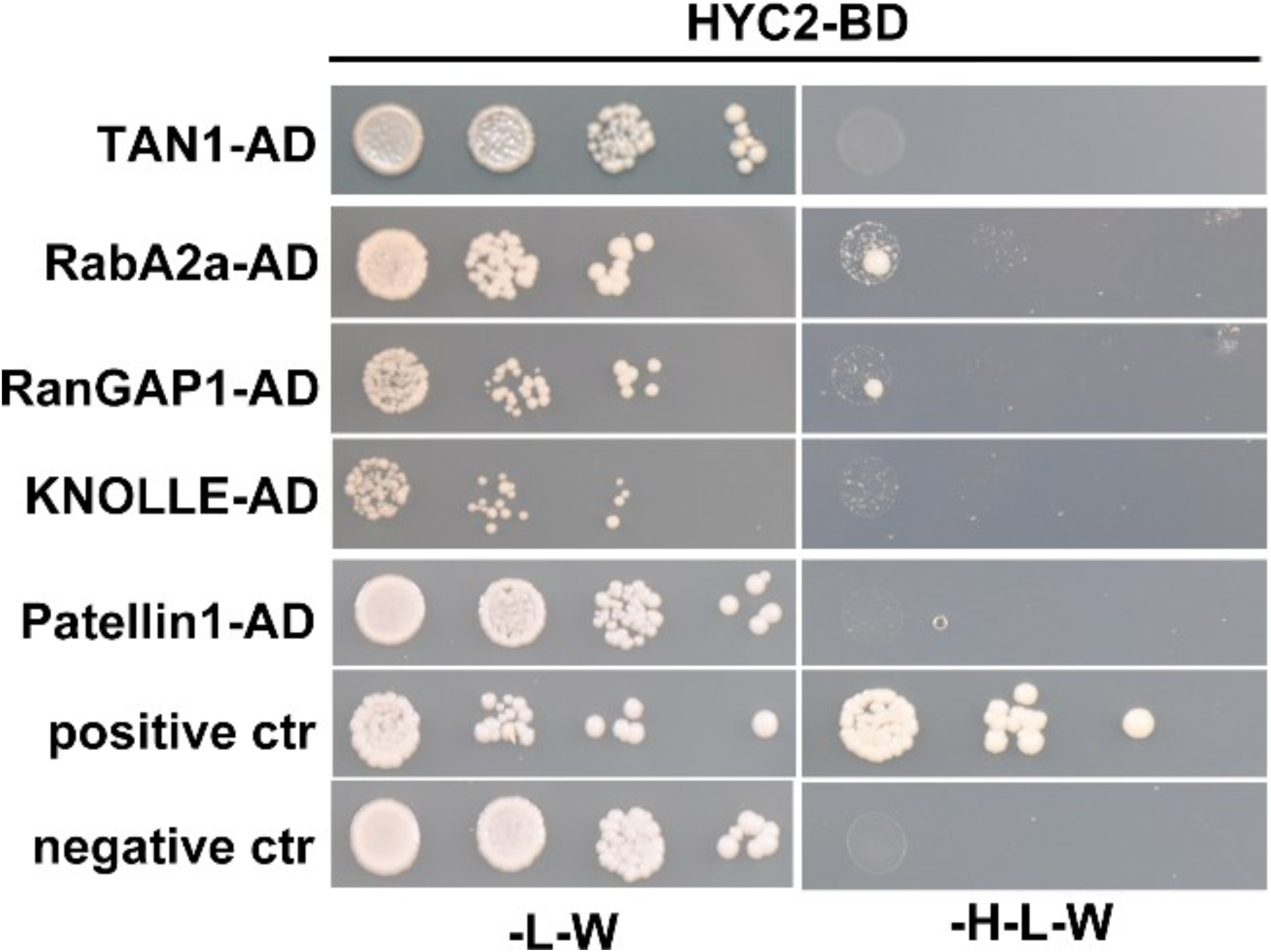
HYC2 protein–protein interactions on Y2H analysis. The protein–protein interactions involving HYC2 and cytokinesis-related proteins were examined by a Y2H assay. Synthetic dropout (SD) nutrient medium, lacking tryptophan (W) and leucine (L), enables yeast cell growth regardless of protein interactions. Additional histidine (H) depletion in the SD medium was used to monitor the interaction of the tested proteins.

**Table S1.** Genotype analysis of self-pollinated Arabidopsis plants with the *hyc2/+* genotype. Over 100 seeds obtained from *hyc2/+* heterozygotes were germinated on a half-strength MS medium. The seedlings were collected for genotyping after 7 days of incubation in a 16-hr light/8-hr dark cycle at 22°C. The segregation ratio of *hyc2/+* progeny was approximately 1:2:0 (WT: HZ: HM), indicating defective embryogenesis (WT, wild-type; HZ, heterozygote; HM, homozygote; *p*>0.05, χ2 test).

| | WT | HZ | HM | Total | Ratio (WT:<br>HZ) | *p* ($\chi^2$ test,<br>WT: HZ=1:2) |
| --- | --- | --- | --- | --- | --- | --- |
| *hyc2-2/+* | 60<br>(31%) | 129<br>(69%) | 0<br>(0%) | 189 | 1:2.15 | 0.64 |
| *hyc2-3/+* | 61<br>(37%) | 104<br>(63%) | 0<br>(0%) | 165 | 1:1.70 | 0.32 |

**Table S2.** Quantitative analysis of cell number and area in Arabidopsis embryos of WT and *hyc2-2/-*. The cell number and area calculation in Arabidopsis embryos involved using images of SCRI-stained cell walls from longitudinal cross-sections. The values were obtained by processing with ImageJ. Asterisks denote *p* < 0.0001 between the two samples (two-tailed, Student *t* test).

|  | WT-globular, n=20 | *hyc2-2*, n=17 | WT-transition, n=18 |
| --- | --- | --- | --- |
| cell number | 23.15 ± 1.98* | 32.47 ± 2.92 | 33.33 ± 2.89 |
| Area (μm<sup>2</sup>) | 1336.47 ± 113.41 | 1376.29 ± 108.08 | 1776.67 ± 125.19* |

\* $p$ value<0.0001, student's $t$ test

**Table S3.** Potential interaction partners of HYC2 in co-immunoprecipitation analysis. Protein extracts from transgenic plants expressing HYC2-2XGFP were incubated with GFP TrapA beads. Precipitated proteins were eluted, trypsin-digested, and identified by liquid chromatography-tandem mass spectrometry (LC-MS/MS). Proteins found in eluents from both HYC2-2XGFP and the wild type were identified as contaminants and excluded from the results. The score represents the value based on the probability that the peptide is a random match to the spectral data. At the same time, coverage indicates the percentage coverage based on the protein’s amino acids.

| Locus | Protein | 1st Repeat |  | 2nd Repeat |  |
| --- | --- | --- | --- | --- | --- |
|  |  | Score | Coverage | Score | Coverage |
| AT1G49340 | ATPI4K ALPHA | 850.38 | 40.80 | 680.33 | 35.90 |
| AT4G28600 | NPGR2 | 280.33 | 37.50 | 290.32 | 38.20 |
| AT2G43040 | NPG1 | 170.33 | 30.00 | 140.30 | 25.00 |
| AT5G64090 | HYC2 | 110.28 | 35.30 | 140.33 | 44.40 |
| AT1G27460 | NPGR1 | 100.31 | 17.40 | 120.26 | 17.30 |
| AT1G49240 | ACT8 | 80.26 | 25.50 | 130.28 | 39.00 |
| AT3G18780 | ACT2 | 80.26 | 25.50 | 130.28 | 39.00 |
| AT5G09810 | ACT7 | 60.24 | 17.80 | 120.26 | 36.60 |
| AT1G05960 | EFOP3 | 40.27 | 6.20 | 40.23 | 4.30 |
| AT3G46520 | ACT12 | 40.15 | 10.90 | 90.22 | 21.50 |
| AT3G12110 | ACT11 | 40.15 | 10.90 | 80.22 | 18.80 |
| AT5G59370 | ACT4 | 40.15 | 10.90 | 90.22 | 21.50 |
| AT2G37620 | ACT1 | 38.15 | 10.90 | 80.19 | 16.70 |
| AT3G53750 | ACT3 | 38.15 | 10.90 | 80.19 | 16.70 |
| AT5G12250 | TUB6 | 30.26 | 12.50 | 50.34 | 13.60 |
| AT5G23860 | TUB8 | 30.26 | 12.50 | 50.34 | 13.60 |
| AT2G29550 | TUB7 | 28.26 | 12.50 | 60.34 | 16.00 |
| AT5G62690 | TUB2 | 28.26 | 13.10 | 80.34 | 22.00 |
| AT5G62700 | TUB3 | 28.26 | 13.10 | 80.34 | 22.00 |
| AT4G20890 | TUB9 | 20.26 | 10.60 | 66.34 | 19.80 |
| AT1G04820 | TUA4 | 20.18 | 6.20 | 50.26 | 17.80 |
| AT1G50010 | TUA2 | 20.18 | 6.20 | 50.26 | 17.80 |
| AT4G14960 | TUA6 | 20.18 | 6.20 | 50.26 | 17.80 |
| AT5G19770 | TUA3 | 20.18 | 6.20 | 30.23 | 9.60 |
| AT5G19780 | TUA5 | 20.18 | 6.20 | 30.23 | 9.60 |
| AT1G75780 | TUB1 | 18.26 | 10.50 | 58.34 | 17.00 |
| AT1G20010 | TUB5 | 18.26 | 10.50 | 58.34 | 16.90 |
| AT5G44340 | TUB4 |  |  | 60.26 | 12.60 |
| AT5G26850 | EFOP1 |  |  | 10.12 | 1.00 |
| AT5G42080 | DRP1A |  |  | 10.15 | 2.10 |

**Table S4.**
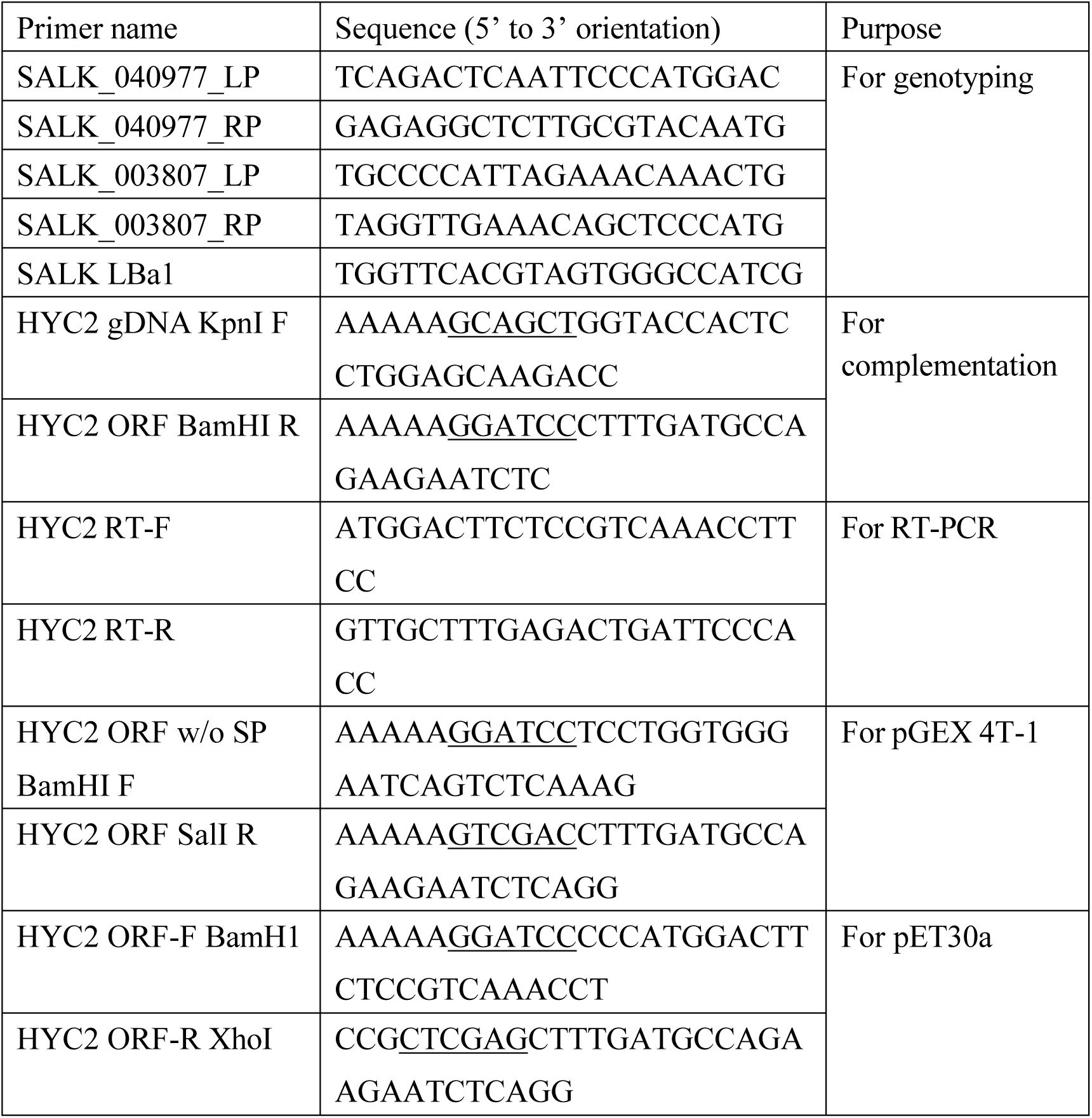
Primer sequences are used for genotyping, cloning, and gene expression; restriction sites within the primers are marked with black underlining.

## SI Methods

### Complementation of the hyc2-2 mutants

To complement the *hyc2-2/-* mutants, the genomic sequence corresponding to locus At5g64090, which includes a 2.5-kb putative promoter and the 1.3-kb coding region, was amplified using Phusion polymerase (Finnzymes, http://www.finnzymes.com) and the *HYC2* gDNA *KpnI F* and *HYC2* ORF *BamHI* R primers. Subsequently, the PCR product was inserted into a modified pPZP221 vector containing two C-terminal GFP sequences followed by the Nopaline synthase (NOS) terminator. Cloning primers are detailed in Table S4. The confirmed construct was introduced into *hyc2-2* heterozygous plants via a floral dip method 1. For antibiotic selection, T1 generation seeds were cultivated on solid half-strength MS medium (Duchefa) supplemented with 1% sucrose, 0.7% phytoagar (Duchefa), and G418 antibiotic at 100 μg/μl. After genotyping, seeds from plants with a *hyc2-2/-* homozygous background carrying *HYC2p:HYC2-2XGFP* were collected for further investigation.

### RNA isolation and RT-PCR analysis

Total RNA was extracted using the Qiagen RNeasy Mini Kit (QIAGEN) and treated with TURBO DNA-free DNase (Ambion) to eliminate genomic DNA contamination. Reverse transcription of RNA was carried out using the M-MLV transcriptase system (Invitrogen) and oligo-dT primers. Primer sequences for *HYC2* expression (*HYC2* RT-F and *HYC1* RT-R) are provided in Table S4.

### Visualization of F-actin in globular embryos

To visualize F-actin in globular embryos, young siliques were fixed in 4% paraformaldehyde in phosphate-buffered saline (PBS), pH 7.4, for 1 h. Subsequently, the buffer was replaced with PBS buffer, pH 7.4, containing 100 nM Alexa 488-phalloidin. After 1 h of incubation, SCRI Renaissance 2200 (SCRI) was added at a 10,000X dilution for 30 min. Embryos were dissected on slides and observed using a Zeiss LSM510 or LSM880 confocal microscope. The skewness, kurtosis, and occupancy of F-actin in the globular embryos were calculated as previously described2. Statistical analysis involved the examination of at least 20 globular embryos from both wild-type and *hyc2-2/-*, analyzed with ImageJ software (http://imagej.nih.gov/ij/).

### Immunofluorescence

Immunofluorescence was performed on whole-mounted roots following the protocol by Sauer and Friml3. Arabidopsis seedlings were mounted on slides with anti-GFP (Abcam, 1:400), anti-tubulin (Sigma, 1:400), and anti-actin (Thermo, 1:100) antibodies. Secondary antibodies for GFP and tubulin were anti-rabbit IgG-FITC and anti-mouse IgG-TRITC, respectively. Before placing coverslips, 100 μM DAPI was used for nuclei staining. Cells were imaged using the Zeiss LSM510 confocal microscope.

## References

Ahn, G., Kim, H., Kim, D.H., Hanh, H., Yoon, Y., Singaram, I., Wijesinghe, K.J., Johnson, K.A., Zhuang, X., Liang, Z., Stahelin, R.V., Jiang, L., Cho, W., Kang, B.H., and Hwang, I. (2017). SH3 Domain-Containing Protein 2 Plays a Crucial Role at the Step of Membrane Tubulation during Cell Plate Formation. Plant Cell 29, 1388–1405.

Audhya, A., Foti, M., and Emr, S.D. (2000). Distinct roles for the yeast phosphatidylinositol 4-kinases, Stt4p and Pik1p, in secretion, cell growth, and organelle membrane dynamics. Mol Biol Cell 11, 2673–2689.

Backues, S.K., Konopka, C.A., McMichael, C.M., and Bednarek, S.Y. (2007). Bridging the divide between cytokinesis and cell expansion. Curr Opin Plant Biol 10, 607–615.

Bannigan, A., Scheible, W.R., Lukowitz, W., Fagerstrom, C., Wadsworth, P., Somerville, C., and Baskin, T.I. (2007). A conserved role for kinesin-5 in plant mitosis. J Cell Sci 120, 2819–2827.

Baskin, J.M., Wu, X., Christiano, R., Oh, M.S., Schauder, C.M., Gazzerro, E., Messa, M., Baldassari, S., Assereto, S., Biancheri, R., Zara, F., Minetti, C., Raimondi, A., Simons, M., Walther, T.C., Reinisch, K.M., and De Camilli, P. (2016). The leukodystrophy protein FAM126A (hyccin) regulates PtdIns(4)P synthesis at the plasma membrane. Nat Cell Biol 18, 132–138.

Benkova, E., Michniewicz, M., Sauer, M., Teichmann, T., Seifertova, D., Jurgens, G., and Friml, J. (2003). Local, efflux-dependent auxin gradients as a common module for plant organ formation. Cell 115, 591–602.

Blilou, I., Xu, J., Wildwater, M., Willemsen, V., Paponov, I., Friml, J., Heidstra, R., Aida, M., Palme, K., and Scheres, B. (2005). The PIN auxin efflux facilitator network controls growth and patterning in Arabidopsis roots. Nature 433, 39–44.

Caillaud, M.C., Lecomte, P., Jammes, F., Quentin, M., Pagnotta, S., Andrio, E., de Almeida Engler, J., Marfaing, N., Gounon, P., Abad, P., and Favery, B. (2008). MAP65-3 microtubule-associated protein is essential for nematode-induced giant cell ontogenesis in Arabidopsis. Plant Cell 20, 423–437.

Criqui, M.C., Parmentier, Y., Derevier, A., Shen, W.H., Dong, A., and Genschik, P. (2000). Cell cycle-dependent proteolysis and ectopic overexpression of cyclin B1 in tobacco BY2 cells. Plant J 24, 763–773.

Dhonukshe, P., Baluska, F., Schlicht, M., Hlavacka, A., Samaj, J., Friml, J., and Gadella, T.W., Jr. (2006). Endocytosis of cell surface material mediates cell plate formation during plant cytokinesis. Dev Cell 10, 137–150.

Drozdetskiy, A., Cole, C., Procter, J., and Barton, G.J. (2015). JPred4: a protein secondary structure prediction server. Nucleic Acids Res 43, W389–394.

Friml, J., Vieten, A., Sauer, M., Weijers, D., Schwarz, H., Hamann, T., Offringa, R., and Jurgens, G. (2003). Efflux-dependent auxin gradients establish the apical-basal axis of Arabidopsis. Nature 426, 147–153.

Gazzerro, E., Baldassari, S., Giacomini, C., Musante, V., Fruscione, F., La Padula, V., Biancheri, R., Scarfi, S., Prada, V., Sotgia, F., Duncan, I.D., Zara, F., Werner, H.B., Lisanti, M.P., Nobbio, L., Corradi, A., and Minetti, C. (2012). Hyccin, the molecule mutated in the leukodystrophy hypomyelination and congenital cataract (HCC), is a neuronal protein. PLoS One 7, e32180.

Hammond, G.R., Fischer, M.J., Anderson, K.E., Holdich, J., Koteci, A., Balla, T., and Irvine, R.F. (2012). PI4P and PI(4,5)P2 are essential but independent lipid determinants of membrane identity. Science 337, 727–730.

Higaki, T., Kutsuna, N., Sano, T., and Hasezawa, S. (2008). Quantitative analysis of changes in actin microfilament contribution to cell plate development in plant cytokinesis. BMC Plant Biol 8, 80.

Higaki, T., Kutsuna, N., Sano, T., Kondo, N., and Hasezawa, S. (2010). Quantification and cluster analysis of actin cytoskeletal structures in plant cells: role of actin bundling in stomatal movement during diurnal cycles in Arabidopsis guard cells. Plant J 61, 156–165.

Ho, C.M., Hotta, T., Guo, F., Roberson, R.W., Lee, Y.R., and Liu, B. (2011). Interaction of antiparallel microtubules in the phragmoplast is mediated by the microtubule-associated protein MAP65-3 in Arabidopsis. Plant Cell 23, 2909–2923.

Jurgens, G., Park, M., Richter, S., Touihri, S., Krause, C., El Kasmi, F., and Mayer, U. (2015). Plant cytokinesis: a tale of membrane traffic and fusion. Biochem Soc Trans 43, 73–78.

Krupnova, T., Sasabe, M., Ghebreghiorghis, L., Gruber, C.W., Hamada, T., Dehmel, V., Strompen, G., Stierhof, Y.D., Lukowitz, W., Kemmerling, B., Machida, Y., Hashimoto, T., Mayer, U., and Jurgens, G. (2009). Microtubule-associated kinase-like protein RUNKEL needed [corrected] for cell plate expansion in Arabidopsis cytokinesis. Curr Biol 19, 518–523.

Lau, S., Slane, D., Herud, O., Kong, J., and Jurgens, G. (2012). Early embryogenesis in flowering plants: setting up the basic body pattern. Annu Rev Plant Biol 63, 483–506.

Lauber, M.H., Waizenegger, I., Steinmann, T., Schwarz, H., Mayer, U., Hwang, I., Lukowitz, W., and Jurgens, G. (1997). The Arabidopsis KNOLLE protein is a cytokinesis-specific syntaxin. J Cell Biol 139, 1485–1493.

Lee, Y.R., Li, Y., and Liu, B. (2007). Two Arabidopsis phragmoplast-associated kinesins play a critical role in cytokinesis during male gametogenesis. Plant Cell 19, 2595–2605.

Li, S., Sun, T., and Ren, H. (2015). The functions of the cytoskeleton and associated proteins during mitosis and cytokinesis in plant cells. Front Plant Sci 6, 282.

Liao, C.Y., and Weijers, D. (2018). A toolkit for studying cellular reorganization during early embryogenesis in Arabidopsis thaliana. Plant J 93, 963–976.

Lin, F., Krishnamoorthy, P., Schubert, V., Hause, G., Heilmann, M., and Heilmann, I. (2019). A dual role for cell plate-associated PI4Kbeta in endocytosis and phragmoplast dynamics during plant somatic cytokinesis. EMBO J 38.

Luptovciak, I., Samakovli, D., Komis, G., and Samaj, J. (2017). KATANIN 1 Is Essential for Embryogenesis and Seed Formation in Arabidopsis. Front Plant Sci 8, 728.

Maeda, K., Sasabe, M., Hanamata, S., Machida, Y., Hasezawa, S., and Higaki, T. (2020). Actin Filament Disruption Alters Phragmoplast Microtubule Dynamics during the Initial Phase of Plant Cytokinesis. Plant Cell Physiol 61, 445–456.

Moller, B., and Weijers, D. (2009). Auxin control of embryo patterning. Cold Spring Harb Perspect Biol 1, a001545.

Mravec, J., Petrasek, J., Li, N., Boeren, S., Karlova, R., Kitakura, S., Parezova, M., Naramoto, S., Nodzynski, T., Dhonukshe, P., Bednarek, S.Y., Zazimalova, E., de Vries, S., and Friml, J. (2011). Cell plate restricted association of DRP1A and PIN proteins is required for cell polarity establishment in Arabidopsis. Curr Biol 21, 1055–1060.

Muller, S., and Livanos, P. (2019). Plant Kinesin-12: Localization Heterogeneity and Functional Implications. Int J Mol Sci 20.

Nakatsu, F., Baskin, J.M., Chung, J., Tanner, L.B., Shui, G., Lee, S.Y., Pirruccello, M., Hao, M., Ingolia, N.T., Wenk, M.R., and De Camilli, P. (2012). PtdIns4P synthesis by PI4KIIIalpha at the plasma membrane and its impact on plasma membrane identity. J Cell Biol 199, 1003–1016.

Noack, L.C., Bayle, V., Armengot, L., Rozier, F., Mamode-Cassim, A., Stevens, F.D., Caillaud, M.C., Munnik, T., Mongrand, S., Pleskot, R., and Jaillais, Y. (2022). A nanodomain-anchored scaffolding complex is required for the function and localization of phosphatidylinositol 4-kinase alpha in plants. Plant Cell 34, 302–332.

Preuss, M.L., Schmitz, A.J., Thole, J.M., Bonner, H.K., Otegui, M.S., and Nielsen, E. (2006). A role for the RabA4b effector protein PI-4Kbeta1 in polarized expansion of root hair cells in Arabidopsis thaliana. J Cell Biol 172, 991–998.

Rasmussen, C.G., Humphries, J.A., and Smith, L.G. (2011). Determination of symmetric and asymmetric division planes in plant cells. Annu Rev Plant Biol 62, 387–409.

Schlereth, A., Moller, B., Liu, W., Kientz, M., Flipse, J., Rademacher, E.H., Schmid, M., Jurgens, G., and Weijers, D. (2010). MONOPTEROS controls embryonic root initiation by regulating a mobile transcription factor. Nature 464, 913–916.

Smertenko, A., Hewitt, S.L., Jacques, C.N., Kacprzyk, R., Liu, Y., Marcec, M.J., Moyo, L., Ogden, A., Oung, H.M., Schmidt, S., and Serrano-Romero, E.A. (2018). Phragmoplast microtubule dynamics - a game of zones. J Cell Sci 131.

Smith, Z.R., and Long, J.A. (2010). Control of Arabidopsis apical-basal embryo polarity by antagonistic transcription factors. Nature 464, 423–426.

Sollner, R., Glasser, G., Wanner, G., Somerville, C.R., Jurgens, G., and Assaad, F.F. (2002). Cytokinesis-defective mutants of Arabidopsis. Plant Physiol 129, 678–690.

Stevenson-Paulik, J., Love, J., and Boss, W.F. (2003). Differential regulation of two Arabidopsis type III phosphatidylinositol 4-kinase isoforms. A regulatory role for the pleckstrin homology domain. Plant Physiol 132, 1053–1064.

Strompen, G., El Kasmi, F., Richter, S., Lukowitz, W., Assaad, F.F., Jurgens, G., and Mayer, U. (2002). The Arabidopsis HINKEL gene encodes a kinesin-related protein involved in cytokinesis and is expressed in a cell cycle-dependent manner. Curr Biol 12, 153–158.

Traverso, M., Assereto, S., Gazzerro, E., Savasta, S., Abdalla, E.M., Rossi, A., Baldassari, S., Fruscione, F., Ruffinazzi, G., Fassad, M.R., El Beheiry, A., Minetti, C., Zara, F., and Biancheri, R. (2013). Novel FAM126A mutations in hypomyelination and congenital cataract disease. Biochem Biophys Res Commun 439, 369–372.

Ugur, S.A., and Tolun, A. (2008). A deletion in DRCTNNB1A associated with hypomyelination and juvenile onset cataract. Eur J Hum Genet 16, 261–264.

Vermeer, J.E., Thole, J.M., Goedhart, J., Nielsen, E., Munnik, T., and Gadella, T.W., Jr. (2009). Imaging phosphatidylinositol 4-phosphate dynamics in living plant cells. Plant J 57, 356–372.

Yao, L., Janmey, P., Frigeri, L.G., Han, W., Fujita, J., Kawakami, Y., Apgar, J.R., and Kawakami, T. (1999). Pleckstrin homology domains interact with filamentous actin. J Biol Chem 274, 19752–19761.

Zara, F., Biancheri, R., Bruno, C., Bordo, L., Assereto, S., Gazzerro, E., Sotgia, F., Wang, X.B., Gianotti, S., Stringara, S., Pedemonte, M., Uziel, G., Rossi, A., Schenone, A., Tortori-Donati, P., van der Knaap, M.S., Lisanti, M.P., and Minetti, C. (2006). Deficiency of hyccin, a newly identified membrane protein, causes hypomyelination and congenital cataract. Nat Genet 38, 1111–1113.

Zhuang, X., Wang, H., Lam, S.K., Gao, C., Wang, X., Cai, Y., and Jiang, L. (2013). A BAR-domain protein SH3P2, which binds to phosphatidylinositol 3-phosphate and ATG8, regulates autophagosome formation in Arabidopsis. Plant Cell 25, 4596–4615.

## References

Clough, S.J. & Bent, A.F. Floral dip: a simplified method for Agrobacterium-mediated transformation of Arabidopsis thaliana. Plant J 16, 735–743 (1998).

Higaki, T., Kutsuna, N., Sano, T., Kondo, N. & Hasezawa, S. Quantification and cluster analysis of actin cytoskeletal structures in plant cells: role of actin bundling in stomatal movement during diurnal cycles in Arabidopsis guard cells. Plant J 61, 156–165 (2010).

Sauer, M. & Friml, J. Immunolocalization of proteins in plants. Methods Mol Biol 655, 253–263 (2010).

